# Osteochondrogenic Transdifferentiation of Vascular Smooth Muscle Cells and Microenvironmental Dynamics in Medial Arterial Calcification

**DOI:** 10.1101/2025.10.28.684711

**Authors:** Yasuhisa Nakao, Tomohisa Sakaue, Jumpei Ito, Mika Hamaguchi, Kayo Takahashi, Masayoshi Kukida, Fumiyasu Seike, Jun Aono, Shigemiki Omiya, Yuji Sakahashi, Morikatsu Yoshida, Masaki Wakabayashi, Manabu Shirai, Kazunori Yasuda, Katsuji Inoue, Shuntaro Ikeda, Osamu Yamaguchi

## Abstract

**Background:** Vascular calcification, particularly medial arterial calcification (MAC), considerably affects cardiovascular mortality. Current treatments are limited because of the unclear molecular mechanisms of MAC. This study aimed to establish an MAC mouse model using O-ring-induced transverse aortic constriction (OTAC) and to identify critical genes, pathways, and cellular interactions involved in MAC formation by combining lineage tracing technology and single-cell RNṣA sequencing (scRNA-seq).

**Methods:** We developed an OTAC mouse model to mimic MAC. Adult C57BL/6J male mice underwent OTAC, and subsequent analyses were performed at various intervals. Histological and immunohistochemical evaluations were conducted to observe changes in vascular smooth muscle cells (VSMCs). Lineage tracing was used to confirm the origin of osteochondrogenic cells in the tunica media. Additionally, scRNA-seq was performed to capture the dynamic changes in VSMCs and other cell types involved in MAC development.

**Results:** Osteochondrogenic cells and the subsequent MAC in OTAC mice were observed using histological evaluations. Immunohistochemistry showed time-dependent decreases in α-SMA expression and increases in SOX9 and RUNX2 expression in VSMCs. Lineage tracing using *Myh11CreERT2;ROSA26-EGFP* mice confirmed that osteochondrogenic cells originated from contractile VSMCs. scRNA-seq confirmed osteochondrogenic transdifferentiation of VSMCs and highlighted dynamic changes in surrounding cells involved in calcification-related and inflammatory processes.

**Conclusion:** The OTAC method enabled a comprehensive understanding of the microenvironmental dynamics during MAC progression at the single-cell level. This powerful model provides a robust platform for future therapeutic intervention research.

**Graphic Abstract:** 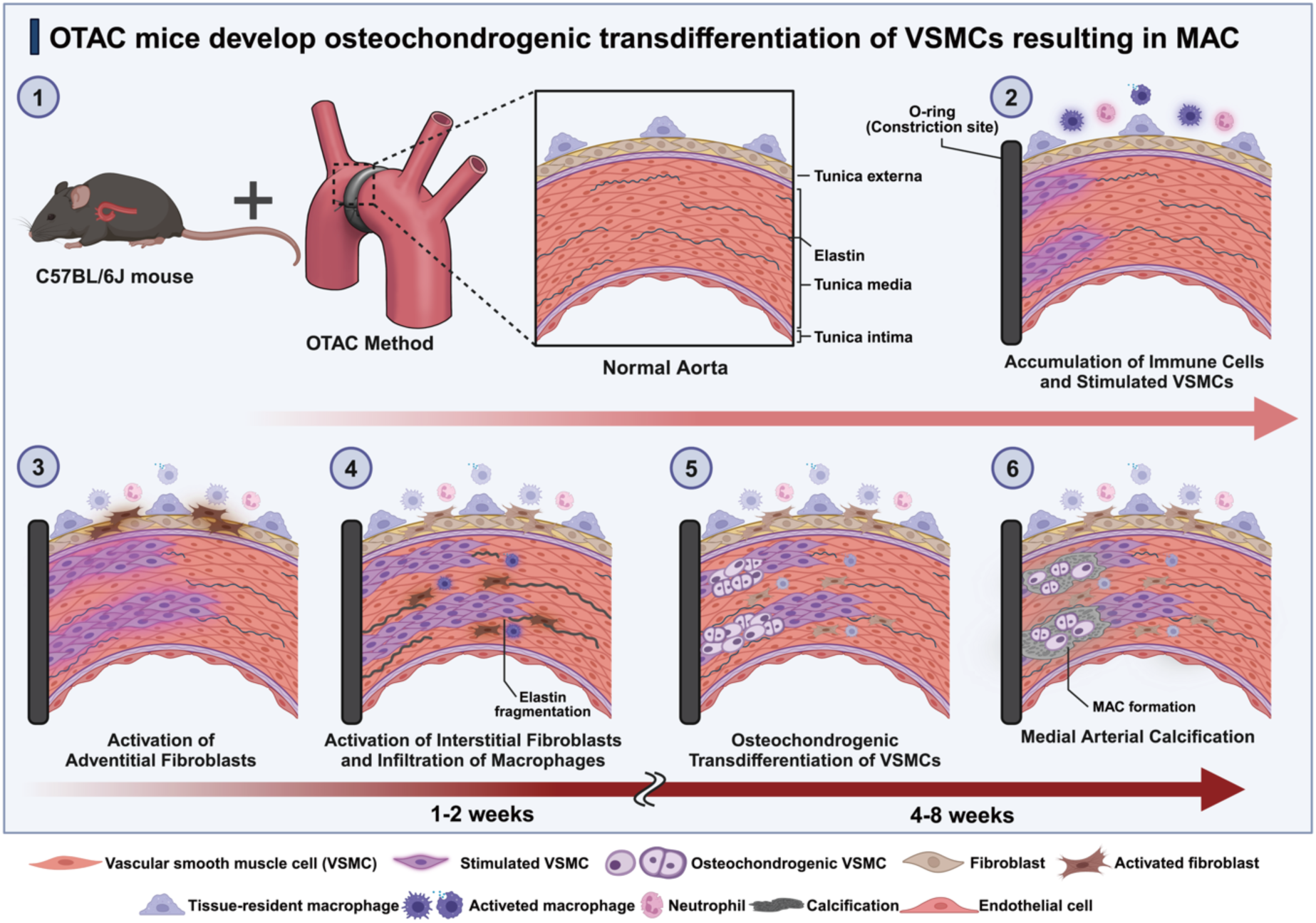

## Introduction

Vascular calcification resulting from hydroxyapatite accumulation considerably contributes to cardiovascular mortality^1,2^. Vascular calcification manifests in two distinct arterial layers: the tunica intima and tunica media^3^. Tunica intima calcification is linked to atherosclerosis, characterized by plaque and cholesterol accumulation within arterial walls^4^. Conversely, medial arterial calcification (MAC) involves calcium deposits in the tunica media and is associated with dyslipidemia, diabetes, chronic kidney disease, and aging^3,5,6^. Despite its connection to severe conditions, treatments for MAC are usually insufficient^7,8^, necessitating comprehensive studies on MAC pathogenesis and potential therapeutic targets^9^. In patients with diabetes, osteochondrogenic transdifferentiation of vascular smooth muscle cells (VSMCs) in tunica media has been reported as a potential cause of MAC^10^, whereas details regarding the specific molecular and cellular interactions underlying MAC remain unknown^3,5^. A critical challenge in MAC research is the lack of an appropriate animal model that accurately replicates human MAC^5,10,11^.

Therefore, to address this issue, we developed a novel MAC mouse model using the O-ring-induced transverse aortic constriction (OTAC) method, which involves applying an O-ring to the transverse aorta^12^. Implanting an O-ring may trigger an outside-in response in the tunica media^13^ and remarkable shear stress on the aorta at its narrowest constriction point, as modeled using Ansys fluent version 18.2 (Ansys Inc., Canonsburg, PA, USA; Supplemental Figure S1A and S1B). We hypothesized that OTAC would stimulate the aorta both internally and externally, leading to MAC development. Thus, this study examined the aortic tissue in OTAC mice to confirm MAC formation, including lineage tracing. Subsequently, we conducted single-cell RNA sequencing (scRNA-seq) analysis to identify the critical genes, signaling pathways, and cellular interactions between VSMCs and surrounding cells for MAC progression.

## Methods

The detailed methods are provided in the Supplemental Methods section. For comprehensive information regarding materials and reagents, please refer to the Major Resources Table.

## Data Availability

Our scRNA-seq and bulk RNA sequencing (RNA-seq) data of mouse aorta are available in the Gene Expression Omnibus under accession numbers GSE268556 and GSE239953, respectively. The scRNA-seq datasets from GSE155468 were used for comparison with human thoracic aortic aneurysms (TAA)^14^. Bulk RNA-seq data from the Sequence Read Archive dataset (PRJNA659049) were used for comparison with the mouse aorta in traditional transverse aortic constriction (TAC) procedures^15^.

## Results

### Calcification with the Appearance of Osteochondrogenic Cells in the Tunica Media of Arteries in OTAC Mice

For histological analysis, aortas from 1-, 2-, 4-, and 8-week post-OTAC and 0-week control mice were evaluated (Figure 1A). The OTAC-induced stenosis region was identified in longitudinal sections (Figure 1B, yellow arrowheads), enabling a detailed examination of adjacent segments. Hematoxylin and eosin staining revealed increased cell density and medial thickness in the proximal and distal regions (Figure 1C). Higher-magnification images (Figure 1D) demonstrated a disorganized arrangement of VSMCs, multilayering, and osteochondrocyte-like cells (green arrowheads) within the media. In addition, mitotic figures were frequently observed among VSMCs (red arrowheads), indicating proliferative activity. Medial thickness peaked at 4-week post-OTAC, and subsequently decreased at 8-week post-OTAC with quantitative analysis (n = 10) confirming a significant increase compared with controls (Figure 1E). An increase in Sirius Red staining was evident in the interstitium of VSMCs during the post-OTAC period (Figure 1C). Additionally, elastin fragmentation was observed in the tunica media (Figure 1D, white arrowheads), and with progressive Alcian blue staining (Figure 1C). Osteochondrocyte-like cells appeared at 4- and 8-week post-OTAC (Figure 1F), which was accompanied by positive von Kossa and Alizarin red S staining beginning at 4 weeks (Figure 1G, and 1H). At 8-week post-OTAC, overt medial calcification was observed in 5 of 10 mice (50%). Representative images from multiple animals are provided in Supplemental Figure S2, confirming reproducibility across specimens. Longitudinal von Kossa staining further demonstrated that calcification was confined to segments immediately adjacent to the O-ring, without spread to distant aortic regions (Supplemental Figure S2A). Notably, early microcalcifications were already detectable at 4 weeks, progressing to larger deposits by 8 weeks (Supplemental Figure S2B). Echocardiography (n = 10) further demonstrated progressive left ventricular hypertrophy (representative images in Supplemental Figure S3A) and reduction in fractional shortening, consistent with increased afterload during the observation period (quantitative analysis in Supplemental Figure S3B).

**Figure 1.**
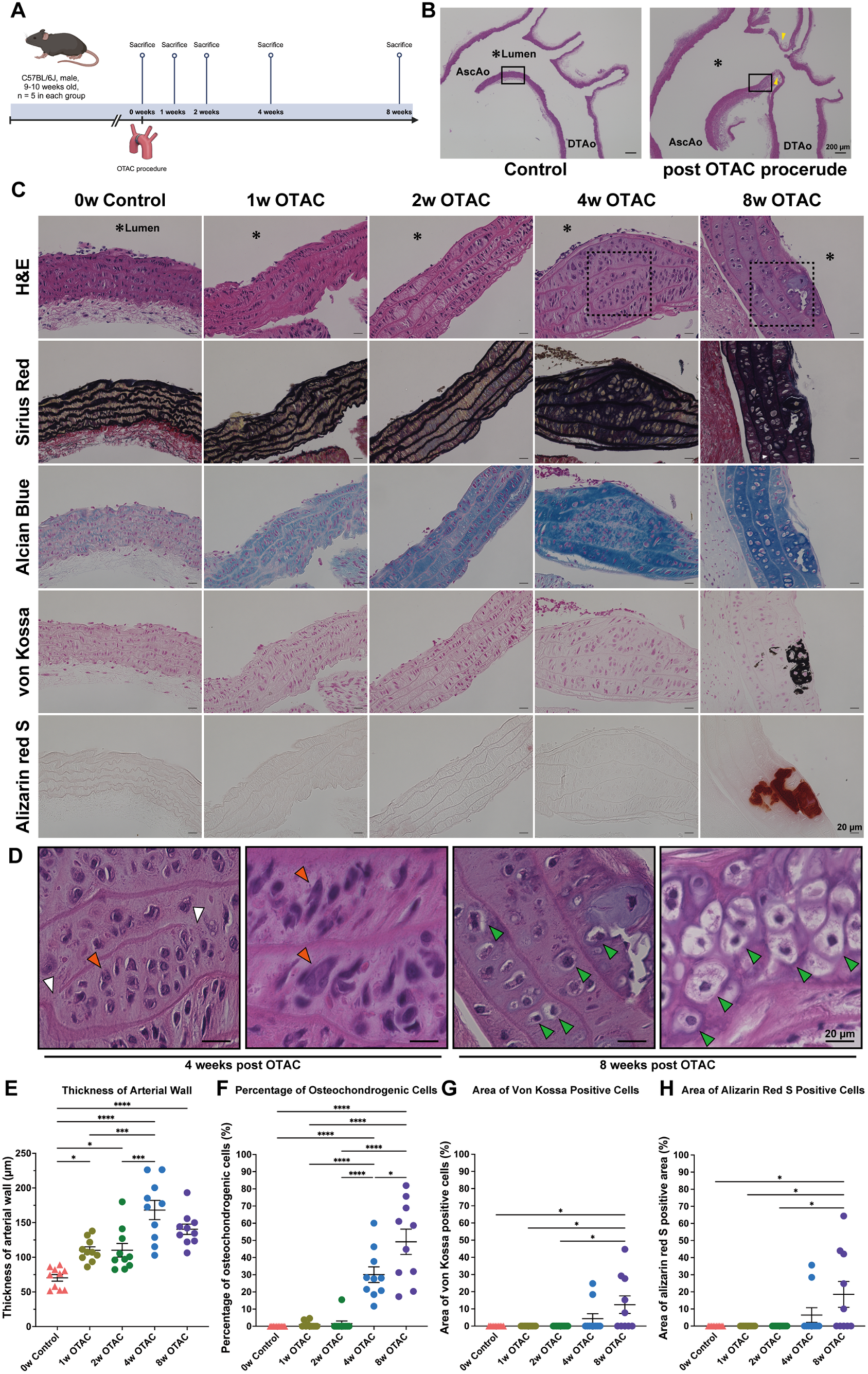
Aortic tissue images and results in OTAC and control mice. **A.** Study overview. C57BL/6J mice (9–10 weeks, weighing 25.5–26.5g) were utilised for the O-ring-induced transverse aortic constriction (OTAC) procedure and control groups (n = 10 per group). **B.** Representative hematoxylin and eosin (HE) staining of the aortic tissue. Scale bar, 200 µm. Yellow arrowheads indicate the constriction site. **C.** Representative aortic tissue images for HE, Sirius Red, Alcian Blue, von Kossa, and Alizarin Red S staining across groups. **D.** Higher-magnification views of the dotted boxed regions in Figure 1C and Supplemental Figure 2B. Arrowheads indicate denucleated cells (green arrowheads), fragmentation of elastin (white arrowheads), and osteochondrogenic cells (red arrowheads). Scale bar, 20 µm. **E.** Quantitative data of the arterial wall thickness. **F**. Quantification of osteochondrogenic cells, expressed as the percentage of osteochondrogenic cells relative to total medial nuclei per section. Each dot represents an individual mouse. **G-H.** Quantification of calcified area by von Kossa (G)/Alizarin red S (H), shown as individual data points per mouse. Histological analysis was performed using n = 10 per group. Data are presented as mean ± standard error of the mean. Group comparisons were performed using one-way analysis of variance with Tukey’s post-hoc test. Significance levels are denoted as *P < 0.05; **P < 0.01; ***P < 0.001; ****P < 0.0001. AscAo, ascending aorta; DTAo, descending thoracic aorta; w, week. The illustration was created with BioRender.com.

### Time-Dependent Changes in α-SMA, SOX9, and RUNX2 Protein Expressions during MAC Formation

Immunohistochemical analysis revealed time-dependent changes in α-SMA, SOX9, and RUNX2 expressions. A marked decrease was found in the levels of contractile protein α-SMA over time (Supplemental Figure S4A). At 4- and 8-week post-OTAC, α-SMA-positive protein levels decreased considerably compared with the baseline control (Supplemental Figure S4B). Conversely, the chondrogenic transcription factor SOX9 and the osteochondrogenic transcription factor RUNX2 expression increased (Supplemental Figure S4A). At 8-week post-OTAC, SOX9 and RUNX2 expression levels were increased compared to those at 0-week control and 1- and 2-week post-OTAC.

### Identification of α-SMA and SOX9 Double-Positive Intermediators in the Tunica Media at OTAC Site

Immunofluorescence staining of α-SMA and SOX9 was performed to evaluate the transdifferentiation of α-SMA-positive VSMCs into osteochondrocyte-like cells. Over time, a decrease in α-SMA-positive VSMCs and a concurrent increase in SOX9-positive VSMCs were observed post-OTAC (Figure 2A). At 4-week post-OTAC, a clear colocalization of α-SMA and SOX9 within VSMCs was detected (Figure 2A and 2B).

**Figure 2.**
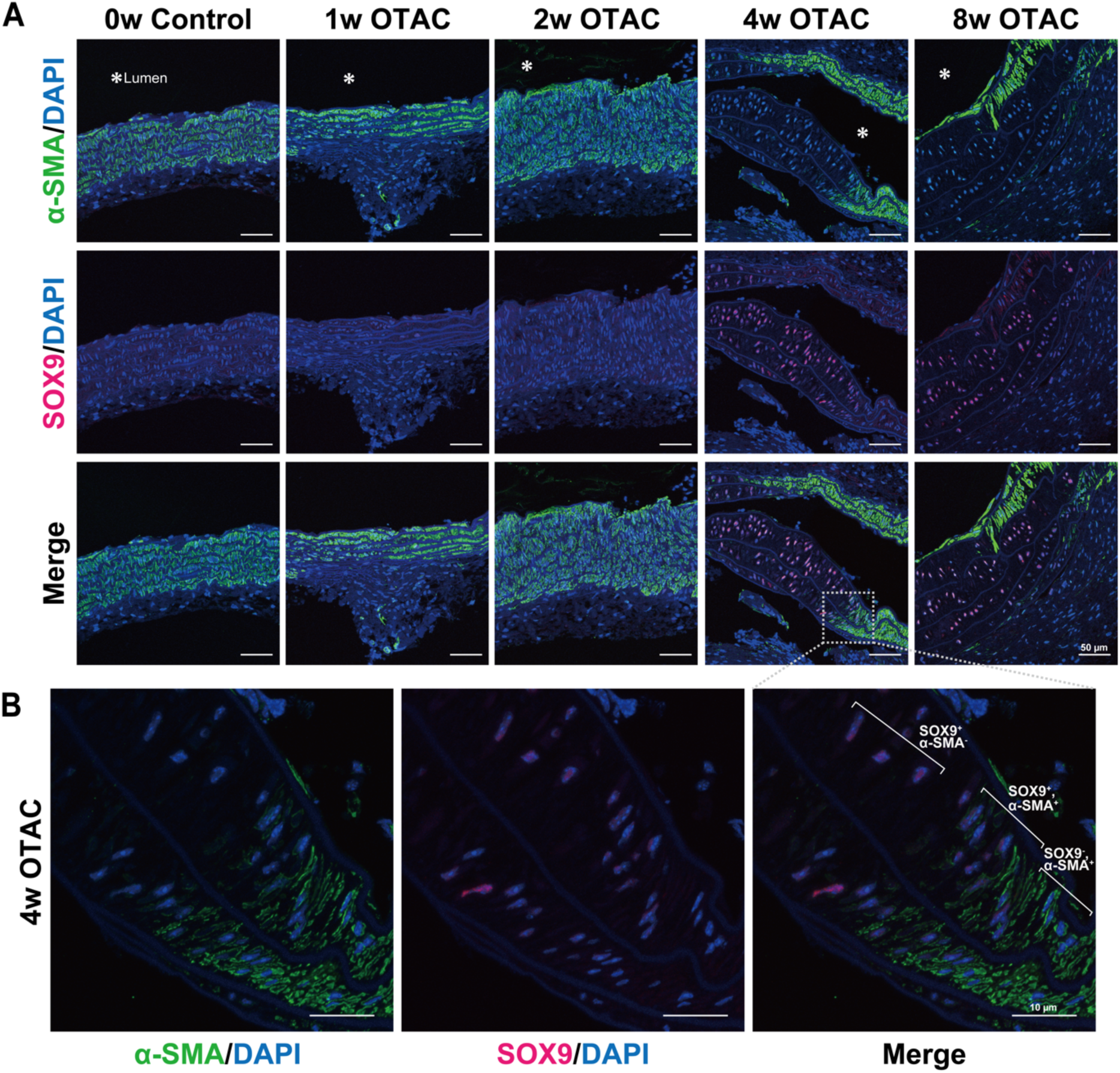
Transition of vascular smooth muscle cells from α-SMA-to SOX9-positive cells. **A.** Representative double immunofluorescence staining of α-smooth muscle actin (α-SMA) and SRY-box transcription factor 9 (SOX9) in aortas from 1-, 2-, 4-, and 8-week post-O-ring-induced transverse aortic constriction (OTAC) and 0-week control mice. The white dotted line demarcates the area detailed in **B.** Scale bar, 50 µm. **B**. Close-up view of the aorta from mice at 4-week post-OTAC. Scale bar, 10 µm. DAPI, 4’,6-diamidino-2-phenylindole; w, week

To complement these protein-level observations, we also examined transcript-level dynamics using bulk RNA-seq (n = 5 per group). K-means clustering stratified genes into six groups (Supplemental Figure S5A), which could be broadly categorized into three temporal patterns. Groups 1 and 3 showed sustained downregulation, including contractile VSMC markers such as *Acta2, Tagln*, *Myh11*, *Cnn1*, and *Myocd*, indicating progressive loss of the contractile phenotype (Supplemental Figure S5B). In addition, regulators such as *Bmp2*, *Smad9*, and *Wnt2b* were also downregulated, suggesting suppression of canonical signaling pathways that normally maintain vascular homeostasis. In contrast, Groups 2, 5, and 6 exhibited early upregulation, encompassing osteochondrogenic regulators such as *Sox9*, *Runx2*, and *Spp1*, consistent with prompt activation of osteochondrogenic programs after OTAC (Supplemental Figure S5B). Early induction of ECM-related genes such as *Fn1* and *Acan*, as well as inflammatory mediators including *Il6* and *Cd68*, further supports a transition toward a pro-calcific and pro-inflammatory milieu. Of note, Group 6 also contained proliferation-associated genes (*Aurkb*, *Cdk1*, *Mki67*, *Mcm2*), pointing to an early burst of cell-cycle activation within the medial layer. Finally, Group 4 displayed late upregulation, including *Col2a1*, *Col10a1*, *Sp7*, and *Prg4*, suggesting a role in subsequent extracellular matrix remodelling and maturation of the osteochondrogenic phenotype (Supplemental Figure S5B).

### Absence of Calcification with Osteochondrogenic Cells in the Tunica Media of Arteries in the Conventional TAC and Its Emergence in the Loose-Tie OTAC Model

To investigate whether MAC formation resulted from shear stress due to aortic constriction or external stimuli from ring placement, we performed both the conventional TAC and loose-tie OTAC procedures (Supplemental Figure S6A). Histological analysis showed the absence of osteochondrogenic VSMCs in the conventional TAC group, while increased intensity of Alcian blue staining was observed (Supplemental Figure S6B). In contrast, VSMCs in mice subjected to loose-tie OTAC differentiated into osteochondrogenic VSMCs and displayed MAC formation (Supplemental Figure S6B).

We indirectly compared our OTAC bulk RNA-seq data with publicly available TAC RNA-seq data from the PRJNA6590434 dataset^15^, aligning the comparison with the similar post-intervention weeks. Using the criteria of fold change > 2.0 and false discovery rate < 0.05, 1,988 upregulated and 772 downregulated differentially expression genes (DEGs) were identified at 2-week post-OTAC mice compared with 0-week control mice. Immune response-related terms in upregulated DEGs and muscle cell-associated terms in downregulated DEGs were enriched in the Gene Ontology (GO) biological process (Supplemental Figure S7A). Gene Set Enrichment Analysis also revealed immune response terms (Supplemental Figure S7B). The top five upregulated and downregulated GO terms in OTAC mice are shown in Supplemental Figure S7C and S7D, respectively. Using equivalent criteria, 798 upregulated and 358 downregulated DEGs were identified at 2-week post-conventional TAC in mice compared with 0-week control mice (Supplemental Figure S7E). Gene set enrichment analysis also revealed immune response- and energy metabolism-related terms (Supplemental Figure S7F). The top five upregulated and downregulated GO terms in TAC mice are shown in Supplemental Figures S7G and S7H. Notably, more DEGs were identified in the OTAC data than in the TAC data, and the downregulated DEGs enriched muscle cell-related terms in the OTAC data.

### Single-cell Profile of Aorta Pre- and Post-OTAC Procedure

To explore dynamic molecular changes in osteochondrogenic transdifferentiation of VSMCs at the single-cell level, we conducted scRNA-seq on ascending to thoracic descending aortic tissues from six male mice at four-time points (pre-OTAC and 1-, 2-, and 4-week post-OTAC) (Figure 3A). After quality control and removal of the doublets, 17,001, 21,353, 18,681, and 5,708 cells remained in the 0-week control, 1-, 2-, and 4-week post-OTAC groups, respectively (Supplemental Table S1). These 62,743 cells were utilized for subsequent analyses.

**Figure 3.**
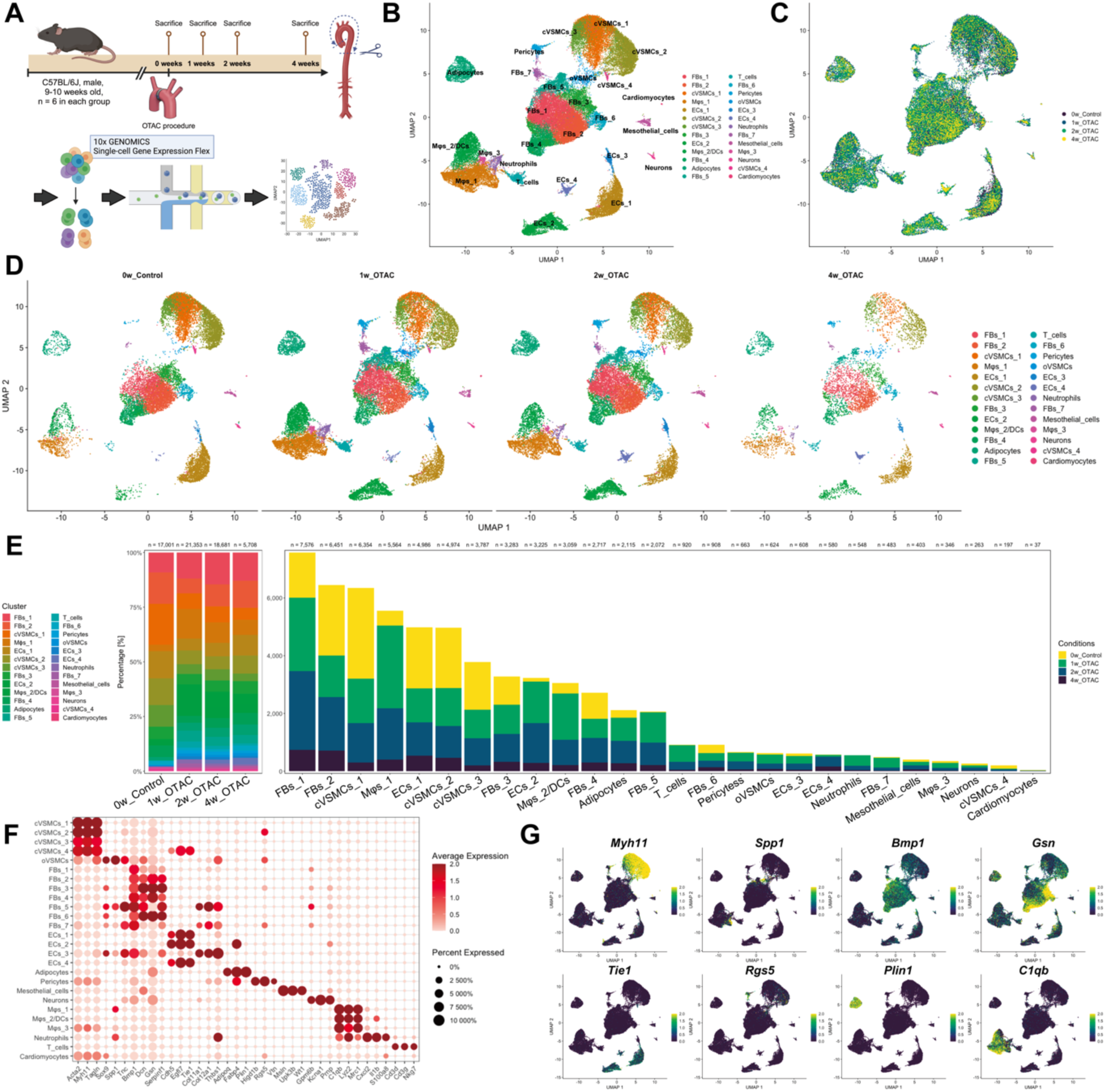
Characterization of all cell types in the single-cell RNA-sequencing data from the aortic tissues of post-OTAC and control mice. **A.** The flow chart of the study, including the study groups and single-cell RNA sequencing (scRNA-seq) platform. **B-C.** Uniform Manifold Approximation and Projection (UMAP) plots of all cells, with each cell color-coded for its associated cell type (**B**) and originating group (**C**). The scRNA-seq datasets were derived from the following groups: 0-week control (0w_Control), 1-week post-O-ring-induced transverse aortic constriction (1w_OTAC), 2-week post-OTAC (2w_OTAC), and 4-week post-OTAC (4w_OTAC). Cell numbers for each group are as follows: 0w_Control (n = 17,001 cells from six mice), 1w_OTAC (n = 21,353 cells from six mice), 2w_OTAC (n = 18,681 cells from six mice), and 4w_OTAC (n = 5,708 cells from six mice). **D.** UMAP plots of all cells divided by the originating group. **E.** Proportion of cell types in each group (**left**). Number of cells in each cell cluster, colored by group (**right**). **F.** Dot plot showing the classic marker genes used to define all cell types. **G.** UMAP plot of all cells colored by the relative expression of classic markers. VSMC, vascular smooth muscle cell; cVSMC, contractile VSMC; oVSMC. osteochondrogenic VSMC; EC, endothelial cell; FB, fibroblast; Mφ, macrophage; DC, dendritic cell. The illustration was created with BioRender.com.

We used 3,000 variable genes per cell to identify cells with similar profiles. Subsequently, we clustered the 26 cell groups using unsupervised clustering and visualized the results employing uniform manifold approximation and projection (UMAP) plots (Figure 3B). Vascular cell types represented in distinct clusters included contractile VSMCs, osteochondrogenic VSMCs, endothelial cells, fibroblasts, macrophages, dendritic cells (DCs), adipocytes, pericytes, T cells, neutrophils, mesothelial cells, cardiomyocytes, and neurons. Overall, fibroblasts constituted the largest population, followed by VSMCs and macrophages, with some clusters exhibiting changes in cell numbers pre- and post-OTAC (Figure 3B-3E). Each cluster was identified using well-known marker genes, and cell type-specific markers were based on the genes with the highest DEGs compared to all other clusters (Figure 3F, 3G, and Supplemental Excel File 1).

### Transdifferentiation of Contractile VSMCs into Osteochondrogenic VSMCs

Among the 26 cell clusters, we identified five VSMC clusters (four contractile VSMCs and one osteochondrogenic VSMCs clusters). In total, 15,936 VSMCs (c_VSMCs_1 to c_VSMCs_4 and o_VSMCs) were categorized into 10 clusters based on their marker genes (Figure 4A and Supplemental Excel File 2). Almost all c_VSMCs clusters exhibited high or similar cell numbers in the control mice compared to the OTAC group (Figure 4A and 4B). However, c_VSMCs_4 cells showed a higher number of cells post-OTAC. In all c_VSMCs clusters, the expression of contraction-related genes, such as *Acta2*, *Myh11*, and *Tagln*, was evident (Figure 4C). While all c_VSMCs clusters highly expressed contraction-related genes, their expression gradually decreased in o_VSMCs (Figure 4C). o_VSMCs were rarely detected in control mice but increased in post-OTAC mice (Figure 4B). We confirmed the expression of osteochondrogenic and vascular calcification-related genes such as *Fbln2*, *Mmp2*, *Igfbp4*, *Spp1*, *Sox9*, *Sdc1*, *Sdc3*, and *Thbs1* (Figure 4C)^3,16–21^. In o_VSMCs, *Spp1* was upregulated in the early post-OTAC period, and *Sox9* was expressed 4 weeks post-OTAC (Figure 4C). Immunofluorescence staining was performed to evaluate the localization of α-SMA and SPP1 in VSMCs. α-SMA- and SPP1-positive cells were detected adjacent to the OTAC site in 1-week post-OTAC aorta but were undetected in controls (Figure 4D). The top 10 GO biological process terms of DEGs identified “Ossification” and “Chondrocyte differentiation” in o_VSMCs. Wnt-related terms reportedly involved in osteochondrogenic transdifferentiation were enriched in c_VSMC_4 (Figure 4E)^21^. *Klf4*, which plays a crucial role in the phenotypic switching of VSMCs^22^, was also identified. To further assess pathway activity at single-cell resolution, we performed AUCell^23^ analysis. UMAP feature plots revealed enrichment of ossification, chondrocyte differentiation, and cartilage development pathways predominantly in oVSMCs, and also in adjacent fibroblast clusters and subsets of contractile VSMCs (Supplemental Figure 8).

**Figure 4.**
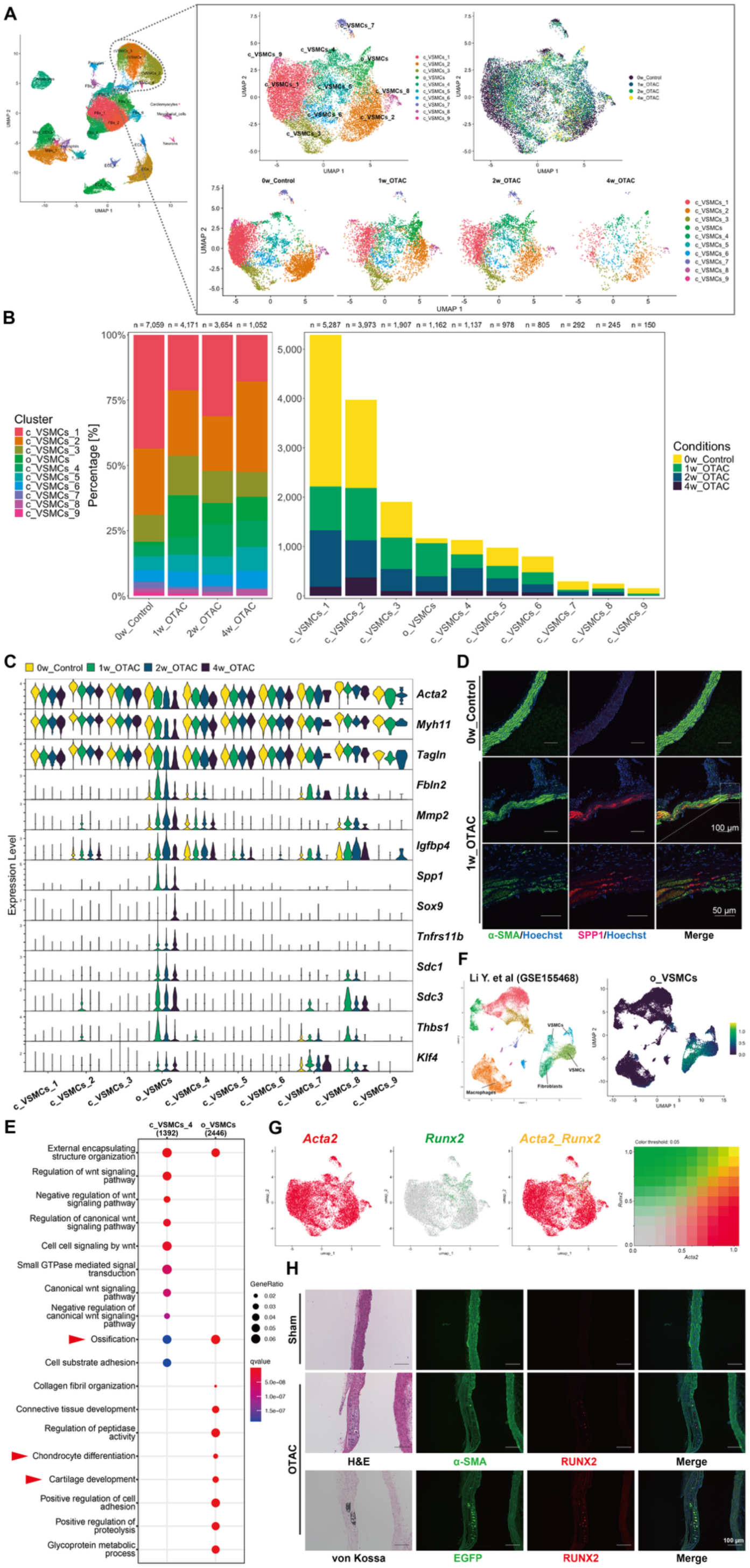
Characterization of subpopulations of vascular smooth muscle cells in OTAC and control mice. **A.** Uniform Manifold Approximation and Projection (UMAP) plot of all vascular smooth muscle cells (VSMCs), colored according to VSMCs sub-clusters (**upper left**), the originating group (**upper right**), and divided by the originating group (**lower**) from 0-week control (0w_Control), 1-week post-O-ring-induced transverse aortic constriction (1w_OTAC), 2-week post-OTAC (2w_OTAC), and 4-week post-OTAC (4w_OTAC). Cell numbers for each group are as follows: 0w_Control (n = 7,059 cells from six mice), 1w_OTAC (n = 4,174 cells from six mice), 2w_OTAC (n = 3,654 cells from six mice), and 4w_OTAC (n = 1,052 cells from six mice). **B.** Proportion of cell types in each group (**left**). Number of cells in each subcluster in each group (**right**). **C.** Violin plot showing markers specific to contractile and osteochondrogenic VSMCs, colored by group. **D.** Representative double immunofluorescence staining of α-SMA and SPP1 in the aortas of 1-week post-OTAC and 0-week control mice. The white dotted line indicates the detailed view. Scale bar, 100 µm. Close-up view of the aorta from 1-week post-OTAC mice. Scale bar, 50 µm. **E.** Top 10 enriched Gene Ontology (GO) biological process terms of the c_VSMCs and o_VSMCs sub-clusters. The red arrowhead indicates GO terms directly associated with vascular calcification. **F.** UMAP visualization from Li et al. scRNA-sequencing data (n = 45,754 cells from eight patients with ascending thoracic aortic aneurysm and three controls) showing the modulation score for o_VSMCs. **G.** UMAP shows the relative expression of *Acta2* (**left**), *Runx2* (**middle**), and both (**right**) in the VSMCs sub-cluster. **H.** Serial sections showing Hematoxylin and Eosin (H&E), immunohistochemical analysis of α-SMA and RUNX2, and adjacent von Kossa staining in sham-operated and OTAC aortas, illustrating the spatial correspondence between calcification and RUNX2-positive regions. Lower panels show colocalization of EGFP and RUNX2 in *Myh11-CreER^T^*^2^*;ROSA26-EGFP* mice, confirming VSMC lineage tracing. Scale bar, 100 µm. α-SMA; α-smooth muscle actin; RUNX2, runt-related transcription factor 2; SPP1, osteopontin; contractile VSMC; oVSMC. osteochondrogenic VSMC.

To detect o_VSMCs-like cells in humans, we analyzed human scRNA-seq data from a TAA study by Li et al^14^ and confirmed the presence of o_VSMCs features-positive cells using a module score calculated from the top 10 DEGs in o_VSMCs (Figure 4F). Based on our scRNA-seq analysis results, some cells within the o_VSMCs cluster were *Acta2*-positive and *Runx2*- or *Sox9*-positive (Figure 4G and Supplemental Figure S9A). To confirm the origin of the osteochondrogenic VSMCs, we performed OTAC in *Myh11-CreER^T^*^2^;*ROSA26-EGFP* mice for VSMCs-specific lineage tracing (Supplemental Figure S9B). Immunofluorescence staining revealed ACTA2-negative and RUNX2- or SOX9-positive cells in aortic tunica media (Figure 4H and Supplemental Figure S9C). To confirm the spatial relationship between these osteochondrogenic cells and calcification, serial von Kossa staining of adjacent sections was performed, demonstrating colocalization of mineral deposition with RUNX2- or SOX9-positive regions (Figure 4H). Conversely, EGFP- and RUNX2- or SOX9-positive cells were detected in aortic tunica media (Figure 4H and Supplemental Figure S9D). To allow direct comparison, sham-operated controls were also analyzed (Figure 4H and Supplemental Figure S9C-S9D). In sham-operated aortas, RUNX2- and SOX9-positive cells were not observed in the medial layer, confirming that the osteochondrogenic changes are specific to the OTAC procedure. Therefore, we determined that RUNX2- and SOX9-positive osteochondrocyte-like cells originated from MYH11-positive contractile VSMCs.

### Dynamics of VSMCs Interaction with Heterogeneous Macrophages During MAC Progression

We classified macrophages and DCs in our scRNA-seq data based on previous reports on atherosclerotic mouse aortas^24–26^. In total, 9,069 macrophages and DCs (Mφs_1, Mφs_2/DCs, and Mφs_3) were categorized into 16 clusters based on cluster-specific marker genes (Figure 5A and Supplemental Excel File 3). Macrophages and DCs were identified as nine (Ma_1 to Ma_9) and five (DCs_1 to DCs_5) subclusters, respectively. One cluster contained *Acta2*- and *Myh11*-positive cells that were not macrophages or DCs but VSMCs. The transition in the number and percentage of each cluster, their specific marker genes, and a corresponding summary are presented in Figure 5B-5D.

**Figure 5.**
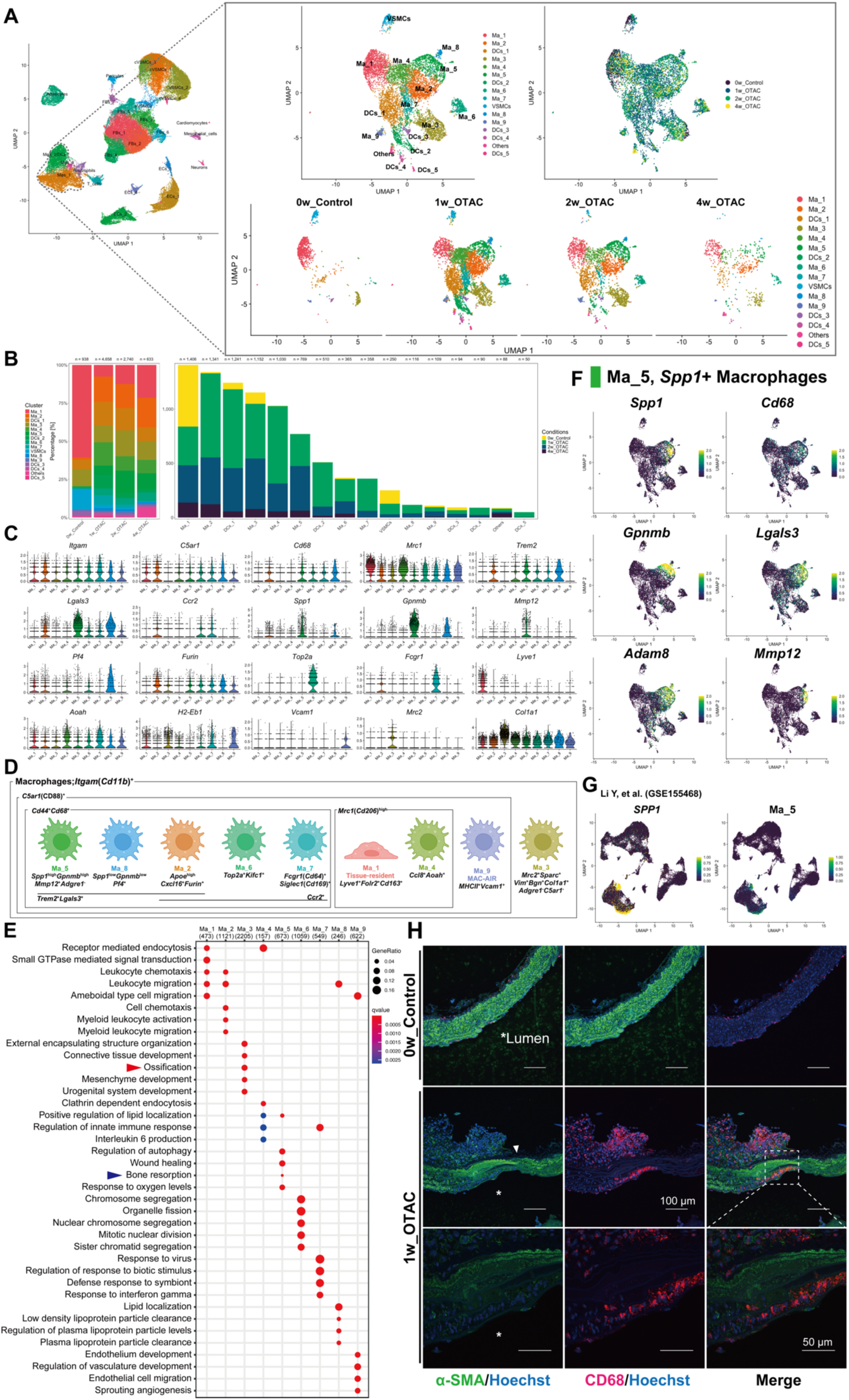
Characterization of macrophage subpopulations in OTAC and control mice. **A.** Uniform Manifold Approximation and Projection (UMAP) plot of all macrophages and dendritic cells (DCs), colored according to macrophages and DCs sub-clusters (**upper left**), the originating group (**upper right**), and divided by the originating group (**lower**) from 0-week control (0w_Control), 1-week post-O-ring-induced transverse aortic constriction (1w_OTAC), 2-week post-OTAC (2w_OTAC), and 4-week post-OTAC (4w_OTAC). Cell numbers for each group are as follows: 0w_Control (n = 938 cells from 6 mice), 1w_OTAC (n = 4,658 cells from six mice), 2w_OTAC (n = 2,740 cells from six mice), and 4w_OTAC (n = 633 cells from six mice). **B.** Proportion of cell types in each group (**left**). Number of cells in each sub-cluster, colored by group (**right**). **C.** Violin plot showing markers specific to macrophages, colored by sub-clusters. **D.** Summary of macrophage subpopulations and their markers from our single-cell RNA-sequencing (scRNA-seq) data. **E.** Top five enriched Gene Ontology (GO) biological process terms of macrophage sub-clusters. Red and blue arrowheads indicate GO terms directly associated with vascular calcification. **F.** UMAP plot of macrophages and DCs subpopulation colored by the relative expression of Ma_5 subcluster-specific markers. **G.** UMAP visualization from Li et al. scRNA-seq data (n = 45,754 cells from eight patients with ascending thoracic aortic aneurysm and three controls) showing *Spp1* and modulation score of Ma_5. **H.** Immunohistochemical analysis of α-SMA and CD68 in 1-week post-OTAC and 0-week control mice. A white arrowhead indicates the constriction site. Scale bar, 100 µm. Close-up view of the aorta from 1-week post-OTAC mice. Scale bar, 50 µm. α-SMA; α-smooth muscle actin; VSMC, vascular smooth muscle cell; Mφ, macrophage. The illustration was created with BioRender.com.

GO enrichment analysis revealed “Bone resorption” terms in Ma_5 (Figure 5E, blue arrowhead), suggesting Ma_5’s involvement in osteochondrogenic processes and aligning with the calcification observed in MAC. Among the macrophage subclusters, Ma_5 was characterized by a high expression of *Spp1*, defining it as *Spp1*-positive macrophages (Figure 5C and 5F). These *Spp1*-positive macrophages also expressed *Gpnmb* and *Mmp12*, which serve as osteoblast differentiation factors (Figure 5F). Using human scRNA-seq data from the TAA study by Li et al.^14^, we confirmed the presence of *SPP1*-positive macrophages and Ma_5-like macrophages from the module score using the top 10 DEGs of Ma_5 (Figure 5G). scRNA-seq analysis confirmed the co-expression of *Spp1* and *Cd68* (Figure 5F). Immunohistochemistry detected CD68-positive cells accumulating in tunica externa and infiltrating tunica media (Figure 5H). Some CD68-positive cells colocalized with α-SMA, indicating the transdifferentiation of macrophage-like VSMCs (Figure 5H)^3,22^. SPP1-positive and α-SMA-negative cells also accumulated surrounding SPP1- and α-SMA-positive cells in the tunica media (Figure 3D).

Other notable macrophage subclusters were provided in this study. Ma_1 and Ma_4 highly expressed *Mrc1*, an M2 macrophage marker. Ma_1, identified as “tissue-resident” macrophages owing to the high expression levels of *Lyve1*, *Cd163*, and *Folr2*, comprising over 60 % of macrophages in control mice, declined post-OTAC but gradually recovered, indicating a dynamic response to vascular injury (Figure 5B). Ma_3 was defined as fibroblast-like macrophages due to the upregulation of *Bgn*, *Col1a1*, *Vim*, *Mrc2*, and *Sparc* and the downregulation of *Adgre1* and *C5ar1* (Figure 5C)^27^. GO enrichment analysis revealed “Ossification” terms in Ma_3 (Figure 5E, red arrowhead). Ma_9 was enriched for the MHC II-encoding transcripts *Vcam1* and *Mrc1*, previously identified as MAC-AIR^25^. *Trem2*-positive macrophages (Ma_2, Ma_5, and Ma_8) were reported in atherosclerotic aorta as “foamy” macrophages^26^, that highly expressed *Trem2* and *Lagls3*.

### Injection of Clodronate Liposome or Neutralization of Neutrophils did not Affect Osteochondrogenic Transdifferentiation of VSMCs

Based on our scRNA-seq results, we hypothesized that the immune response, particularly *Spp1*-positive macrophages and neutrophils (Figure 3F and 3G), play a role in the osteochondrogenic transdifferentiation of VSMCs. Effective macrophage depletion after clodronate liposome treatment in OTAC mice was verified using flow cytometry, focusing on F4/80/CD11c double-positive cells (Supplemental Figure S10A-S10B). However, histological evaluations conducted after the 4-week post-OTAC revealed the presence of osteochondrogenic VSMCs in both clodronate liposome- and control liposome-treated aortas (Supplemental Figure S10C).

We also administered an intraperitoneal injection of Ly6g neutralizing antibodies (nAb) and isotype control antibodies (cAb) to OTAC mice to examine the potential influence of Spp1-positive neutrophils on MAC development. Effective neutrophil removal was verified by flow cytometry, focusing on CD11b/Ly6g double-positive cells (Supplemental Figure S10D-S10E). After the 4-week post-OTAC intervention, histological evaluations revealed osteochondrogenic VSMCs in the aortas of both nAb- and cAb-treated OTAC mice (Supplemental Figure S10F). Therefore, the removal of macrophages and neutrophils alone did not inhibit transdifferentiation into osteochondrogenic VSMCs.

### Activated Fibroblasts Related to Human TAA were Identified in the Aorta of OTAC Mice

In total, 23,494 fibroblasts (FBs_1 to FBs_7) were classified into 11 clusters based on cluster-specific marker genes (Supplemental Figure S11A and Supplemental Excel File 4). Fibroblasts were identified as 11 sub-clusters (Fb_1 to Fb_11). Fb_1 comprised > 60 % of the fibroblasts in control mice and decreased in post-OTAC mice (Supplemental Figure S11B). Common fibroblast markers, such as *Col1a1*, *Col1a2*, *Fbln1*, and *Loxl1,* were expressed in Fb_1 (Supplemental Figure S11C). Fb_2, Fb_3, Fb_8, and Fb_9 were markedly increased in post-OTAC mice (Supplemental Figure S11B). GO enrichment analysis revealed “Ossification” terms in Fb_3 and Fb_4 (Supplemental Figure S11D). Fb_3 was identified as activated fibroblasts characterized by the expression of *Postn*. Fb_3 also expressed several genes associated with tendon and cartilage development, including *Tnc*, *Runx1*, and *Col12a1* (Supplemental Figure S11C). Previously, Klarin et al. reported that *COL6A3* and *FBN1* were risk genes in TAA from a phenome-wide association study, and this was validated in fibroblasts using scRNA-seq data of human TAA obtained from the study by Li et al.^14,28^ Our scRNA-seq data also showed the expression of *Col6a3* and *Fbn1*, mainly in Fb_3 (Supplemental Figure S11C and S11F). Furthermore, *COL16A1* was reported as a candidate gene in TAA with Marfan syndrome in a genome-wide association study^29^. *COL16A1* was also expressed mainly in the fibroblasts of patients with TAA (Supplemental Figure S11G). In Fb_3 cells, we confirmed the co-expression of *Postn* and *Col16a1* in our scRNA-seq data (Supplemental Figure S11C and S11H).

### Cell–Cell Communication Driving Osteochondrogenic VSMC Transdifferentiation

Application of LIANA^30^ with CellPhoneDB^31^ identified extensive intercellular communication across aortic cell populations in the OTAC model (Figure 6A). Category-level network visualization confirmed that fibroblasts and endothelial cells were the predominant signal senders, while VSMCs consistently emerged as dominant receivers of diverse inputs (Figure 6B). Among the calcification-related pathways, INTEGRIN-mediated inputs, particularly driven by SPP1 as well as COL1A1, COL1A2, and FN1, were prominently enriched across fibroblasts, macrophages, and stromal populations (Figure 6C). Pathway-restricted visualization of the Top 30 VSMC-directed interactions highlighted fibroblast-derived ECM-Integrin signaling, including COL3A1-ITGA10/ITGB1, which were selectively augmented after OTAC (Figure 6D). Together, these findings position extracellular matrix signaling as convergent and central pathways orchestrating osteochondrogenic transitions in VSMCs within the pro-calcific microenvironment.

**Figure 6.**
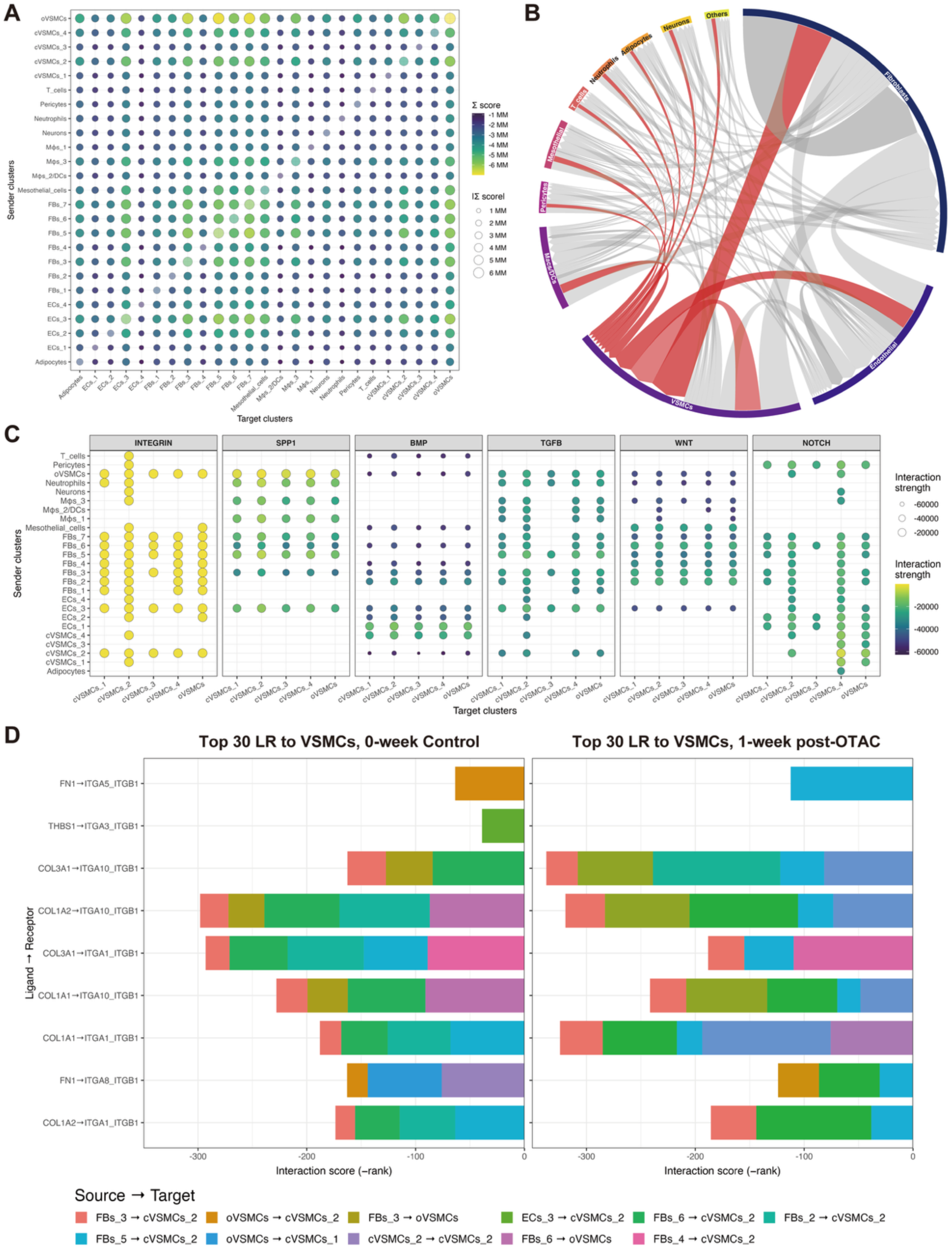
Cell–cell communication dynamics in OTAC model. **A.** Bubble plot summarizing all cluster-to-cluster interactions across the dataset. Each dot represents the summed interaction score (Σ score) between sender (rows) and target (columns) cell clusters. Dot size indicates the absolute magnitude of the total interaction, and dot color encodes the signed interaction score (viridis scale). **B.** Chord diagram summarizing inter-cluster communication at the at the cell type level [e.g., fibroblasts, endothelial cells, vascular smooth muscle cells (VSMCs), etc.]. Link width is proportional to the total interaction strength. Links directed toward VSMCs are highlighted in red, emphasizing their central role as major targets of intercellular signaling. **C.** Calcification-related signaling pathways toward VSMCs. Bubble plots show the top ligand–receptor interactions categorized by pathway (Integrin, SPP1, BMP, TGFB, WNT, NOTCH). Dot size and color represent interaction strength, revealing pathway-specific communication patterns from distinct sender populations. **D.** Top 30 ligand– receptor pairs directed toward VSMCs at baseline (0-week, left) and 1-week post-OTAC (right). Horizontal bar length reflects interaction score (-rank). Colors indicate the contributing source→target cluster combinations. cVSMC, contractile VSMC; EC, endothelial cell; FB, fibroblast; Mφ, macrophage; DC, dendritic cell.

## Discussion

Our study introduced a novel MAC mouse model and identified genes and signaling pathways associated with MAC. The key findings are as follows: (1) MAC was evident in both the proximal and distal segments of the arterial tunica media affected by OTAC-induced stenosis. (2) Osteochondrogenic transdifferentiation occurs directly from VSMCs before calcification. (3) The OTAC procedure stimulates contractile VSMCs and surrounding cells, creating a microenvironment that facilitates osteochondrogenic transdifferentiation of VSMCs. OTAC induced the accumulation of inflammatory cells at the perivascular site, which primarily activated macrophages. Some of these activated macrophages infiltrated the tunica media, activating perivascular and interstitial fibroblasts. Furthermore, changes in the cellular environment surrounding VSMCs influence the osteochondrogenic transdifferentiation of VSMCs, subsequently contributing to MAC formation (Graphical Abstract).

The establishment of a MAC mouse model using the OTAC procedure represents a remarkable strength of our study. This model replicates the pathophysiology of human osteochondrogenic transdifferentiation of VSMCs and MAC^10^, which is crucial for understanding disease processes. MAC was observed between 4- and 8-week post-OTAC, with incidents increasing weekly (Figure 1C, G-H). Traditional models, from *in vitro* cell systems to *in vivo* animal frameworks, have been utilized to study vascular calcification mechanisms^11^, each with different benefits and constraints. For instance, *in vivo* rodent models mimic systemic physiological states; however, calcification patterns largely depend on the inducing agents used. Dietary manipulation models, such as high-fat or cholesterol intake, primarily exhibit intimal calcification, which inadequately captures the MAC phenotype^5^. Therefore, our model emulates human MAC and fills a crucial gap in current research.

The OTAC model validated osteochondrogenic transdifferentiation of VSMCs. A considerable division and disarray of VSMCs in the tunica media were observed post-OTAC, with these cells assuming an osteochondrocyte-like identity. Immunostaining and immunofluorescence revealed a decline in α-SMA-positive VSMCs, reaching a low level between 4 and 8 weeks post-OTAC, while SOX9- and RUNX2-positive cells increased (Figure 2A-B and Supplemental Figure 4A-B). At 4-week post-OTAC, colocalization of α-SMA- and SOX9-positive VSMCs was observed (Figure 2B). Lineage tracing in *Myh11-CreER^T^*^2^;*ROSA26-EGFP* mice with OTAC confirmed the transdifferentiation of MYH11-positive contractile VSMCs into SOX9- and RUNX2-positive osteochondrogenic VSMCs, suggesting that osteochondrogenic VSMCs originated from contractile VSMCs (Figure 4H and Supplemental Figure 9C-D).

Historically, VSMCs have been known for their contractile phenotype, slow proliferation rate, and responsiveness to specific signals^3^. VSMCs produce various contractile proteins, including α-SMA, MYH11, and TAGLN. However, VSMCs exhibit remarkable phenotypic plasticity^32^ and can switch from a contractile to an osteochondrogenic phenotype under certain stimuli. This shift is associated with the formation of calcifying vesicles, the downregulation of mineralization inhibitors, and the production of a matrix prone to calcification. While VSMCs’ plasticity is widely accepted, the diversity of phenotypes they can adopt and their contribution to vascular calcification remains contentious^3,23,33^. Our study shows that VSMCs can transdifferentiate into osteochondrocyte-like cells, contributing to MAC and enriching our understanding of the cellular mechanisms. Although our murine phenotype may not encompass all forms of human MAC, osteochondrogenic cells have been described in human vascular calcification, particularly in diabetic arteries^10^. More recently, SOX9 has been implicated as a key regulator of vascular aging and matrix remodeling^34^, and osteochondrogenic transcription factors have been shown to exhibit distinct expression patterns in human arterial calcification^35^. Together, these reports support the translational relevance of both osteogenic and chondrogenic programs in human arteries, highlighting the applicability of our findings from the OTAC model. Direct evidence of regulatory divergence between intimal and medial calcification will require chromatin accessibility profiling such as scATAC-seq, which remains scarce for medial calcification. We plan to pursue such integrative multi-omics approaches in future work.

We hypothesized that the inflammation and shear stress induced by OTAC could trigger MAC. Histological analysis suggests that inflammation, shear stress, or both might drive this transdifferentiation. To isolate the effects of shear stress, we used a larger O-ring to mitigate shear stress and a traditional TAC to assess shear stress alone (Supplemental Figure S6). Osteochondrogenic transdifferentiation and MAC were observed in the loose-tie OTAC group rather than in the conventional TAC group, highlighting the role of the O-ring in facilitating these processes. Our analyses also demonstrated that these osteochondrogenic changes were regionally restricted to vascular segments adjacent to the O-ring, while remote territories such as the ascending and descending aorta showed little or no alteration. This supports the view that local perivascular stimuli, rather than global hemodynamic stress, are the primary drivers of medial calcification in this model. Previous studies using the TAC approach have highlighted an enlargement in aortic diameter and thickening of its wall^15^, as observed in our study. We also observed some regions of heightened intensity with alcian blue staining in the TAC aorta. RNA-seq analysis showed more DEGs in the OTAC model than in the TAC model, with upregulated inflammation-related DEGs and downregulated contractile VSMC-related DEGs observed in the OTAC model (Supplemental Figure S7). Therefore, while TAC also tends to promote osteochondrogenic differentiation in VSMCs, OTAC induces obvious osteochondrogenic differentiation by eliciting a robust inflammatory response.

The “outside-in” hypothesis posits that aortic wall remodeling involves enhanced activation and migration of fibroblasts and myofibroblasts, as well as increased neovascularization, leukocyte infiltration, and high levels of reactive oxygen species^13,36,37^. These processes exacerbate the recruitment and infiltration of inflammatory cells into arterial walls. The O-ring in our OTAC procedure likely induces this "outside-in" reaction, leading to the osteochondrogenic transdifferentiation of VSMCs and subsequent MAC. Additionally, the direct effects of the O-ring coming into contact with the adjacent VSMCs were considered remarkable. To corroborate the “outside-in” reaction, we performed a scRNA-seq time-series analysis. Contractile VSMCs were present in both post-OTAC and normal aortas but considerably decreased in the post-OTAC aorta (Figure 4A and 4B), indicating that VSMCs were less capable of contracting and transdifferentiating into other phenotypes. We confirmed that osteochondrogenic VSMCs expressed genes involved in MAC formation, such as *Spp1*, *Sox9,* and *Runx2* in post-OTAC mice (Figure 4C and 4D)^3,17^. Although scRNA-seq indicated a relative reduction in osteochondrogenic VSMCs at 4 weeks post-OTAC, this finding contrasts with the abundant osteochondrogenic cells observed in histological sections. This discrepancy is most likely attributable to enzymatic dissociation bias, as mineralized medial regions are difficult to liberate as viable single cells, leading to underrepresentation in scRNA-seq datasets. Supporting the histological results, bulk RNA-seq (n = 5 per group), without single-cell dissociation, demonstrated increased expression of *Runx2*, *Sox9*, and *Spp1* at the same timepoint (Supplemental Figure S5). These complementary data indicate that the decline in scRNA-seq reflects technical limitations rather than a true biological loss of osteochondrogenic VSMCs. cVSMCs_4 subcluster was enriched in the Wnt signaling pathway, contributing to VSMC phenotype regulation in vascular calcification (Figure 4E)^21,38^. Transdifferentiating VSCMs to CD68- and α-SMA-positive macrophage-like VSMCs were also identified (Figure 5H)^3,22^. Some VSMC-like cells were also detected within macrophage/DC clusters. We interpret this as a limitation of the clustering algorithm, whereby partial transcriptomic overlap between VSMCs and macrophages leads to mis-assignment, rather than evidence of bona fide macrophage-like VSMCs. Future spatial or multi-omic analyses will be needed to more clearly delineate these populations. We observed notable phenotypic changes in macrophages and fibroblasts, with *Spp1*-positive macrophages^39^ and neutrophils^40^, endothelial cells, and activated fibroblasts signaling to neighboring cells (Figure 5 and Supplemental Figure S11). AUCell results suggest that not only oVSMCs but also perivascular fibroblasts and subsets of contractile VSMCs may participate in establishing a calcification-prone microenvironment, reinforcing the concept that MAC development is orchestrated by both VSMCs and neighboring stromal cells (Supplemental Figure S8). In particular, macrophages were diverse and heterogeneous post-OTAC, accumulating in the tunica externa and infiltrating around osteochondrogenic VSMCs in tunica media as SPP1-positive macrophages (Figure 4D and 5H). Changes in the surrounding cells centered around osteochondrogenic VSMCs were also observed in arterial remodeling disease, such as human AAA. Nevertheless, direct validation using calcified human aortic tissues remains limited, and future studies with such samples will be critical to fully establish the translational relevance of our findings. To systematically evaluate intercellular communications driving vascular calcification, we applied the LIANA^30^ framework with CellPhoneDB^31^ as the primary resource (Figure 6). This analysis consistently identified extracellular matrix–integrin signaling as the dominant axis directing signals toward VSMCs, with SPP1, COL1A1, COL1A2, and FN1 serving as major upstream ligands from fibroblasts, endothelial cells, and stromal populations (Figure 6A-C). Pathway-restricted visualization of the top VSMC-directed interactions further emphasized fibroblast-derived ECM–integrin inputs, particularly COL3A1–ITGA10/ITGB1 and COL1A1–ITGA1/ITGB1, which were selectively augmented after OTAC (Figure 6D). These findings extend prior observations of SPP1–integrin signaling in vascular calcification by providing a comprehensive, network-level ranking of intercellular pathways, and position integrin-mediated communication as a convergent and central mechanism orchestrating osteochondrogenic transitions of VSMCs *in vivo*. In line with a cooperative mechanism, depletion of macrophages (clodronate) and neutralization of neutrophils (anti-Ly6g) alone did not prevent osteochondrogenic transdifferentiation of VSMCs and medial calcification (Supplemental Figure S10). This indicates that no single cell type is exclusively responsible for the transdifferentiation into osteochondrogenic VSMCs. Together, these findings indicate that OTAC activates both VSMCs and surrounding stromal/immune cells, which cooperatively drive osteochondrogenic transdifferentiation and MAC formation.

Our study has some limitations. Although the MAC mouse model is a good representation of human MAC, species differences may influence disease trajectory and intervention response. The inflammatory reaction induced by the O-ring may differ from human periaortic inflammation and the subsequent calcification processes. Our focus on a single mouse strain may not fully capture the human genetic heterogeneity. Additionally, the presence of osteochondrogenic cells has been confirmed in human atherosclerotic lesions^31^ and MAC^10^; however, their direct association with MAC formation remains unclear. Therefore, further studies, such as spatial transcriptomics, are required to elucidate this relationship. The significance of the perivascular tissue necessitates analyses that preserve this undisturbed tissue. Furthermore, the reduction in cell numbers observed in the scRNA-seq data at 4-week post-OTAC likely reflects the presence of calcified tissue, necessitating a substantial increase in sample numbers for comparability. Moreover, while our transcriptomic analyses identified dynamic expression changes in both known and putative regulators of medial calcification, we did not conduct *in vitro* functional validation in the present study. Conventional VSMC cultures often fail to reproduce the phenotype of contractile VSMCs *in vivo*. Thus, future mechanistic studies will be better addressed *in vivo* using the OTAC model as a platform.

In conclusion, we successfully developed a MAC model mouse allows for reliable tracking of the calcification process and revealed critical insights into the transdifferentiation of VSMCs with microenvironmental dynamics. By enabling detailed studies of the calcification process, it paves the way for the development of targeted molecular treatments.

## Author Contribution Statement

Y.N., T.S., M.K., J.A., and O.Y.: conceptualization; Y.N., T.S., J.I., M.H., K.T., and O.Y.; methodology; Y.N., T.S., J.I., K.T., F.S., and K.Y.: investigation; Y.N., T.S., J.I., K.T., F.S., and K.Y.: visualization; T.S., S.O., and O.Y.: project administration and supervision; Y.N.: writing—original draft;:T.S., J.I., Y.S., M.Y., M.W., M.S., K.I., S.I., and O.Y.: writing— review and editing.

## Acknowledgments

We are grateful to Makiko Takahashi, Isuzu Ikeuchi, and Naohito Tokunaga from the Division of Analytical Biomedicine and the Division of Laboratory Medical Research Support at the Advanced Research Support Center (ADRES), Ehime University, for their professional assistance. We would like to thank Editage [http://www.editage.com] for editing and reviewing this manuscript for English language.

## Supplemental Materials

### Supplemental Methods

Figures S1–S11

Tables S1

Excel File 1-4

References^12,15,23,30–31,41–56^

## Non-standard Abbreviations and Acronyms

cAb: Control antibodies
DCs: Dendritic cells
DEGs: Differentially expressed genes
FC: Fold change
FDR: False discovery rate
GO: Gene Ontology
MAC: Medial arterial calcification
nAb Ly6g: neutralizing antibodies
OTAC: O-ring-induced transverse aortic constriction
RNA-: seq RNA sequencing
scRNA-: seq Single-cell RNA sequencing
TAA: Thoracic aortic aneurysms
TAC: Transverse aortic constriction
VSMCs: Vascular smooth muscle cells

## Highlights

- MAC significantly impacts cardiovascular mortality due to its unclear molecular mechanisms and lack of treatment.
- We provide the first evidence regarding novel MAC model with osteochondrogenic VSMCs in transverse aorta.
- With high reproducibility, quantitative precision, and comprehensive understanding of pathology, our model enables analysis at the single-cell level.
- The insights from our novel model could accelerate the development of direct treatment for MAC.

