## Supplemental_materials for "Osteochondrogenic Transdifferentiation of Vascular Smooth Muscle Cells and Microenvironmental Dynamics in Medial Arterial Calcification"


Yasuhisa Nakao et al.

Corresponding Authors:

Tomohisa Sakaue,

Osamu Yamaguchi,


**The PDF file includes:**

Supplemental Methods**,** Supplemental Figures S1-S11, and Supplemental Tables S1

### Supplemental Methods

#### Ethics

All mouse experiments were approved by the Ehime University Animal Research Committee (approval no.05TA53-1,16) and the National Cerebral and Cardiovascular Center Committee (approval no.22109), and its protocol conformed to the institutional guidelines.

#### Experimental Animals

Adult C57BL/6J male mice, weighing 26.0 ± 0.5 g at 9–10 weeks, were procured from CLEA Japan, Inc., Tokyo, Japan. Mice were categorized into the OTAC operation or control groups. A detailed description of the OTAC procedure has been outlined elsewhere[12]. Briefly, the mice were initially anesthetized using a mixture of medetomidine, midazolam, and butorphanol (0.3, 4.0, and 5.0 mg/kg, respectively) with intraperitoneal administration. The mice underwent aortic constriction using nitrile rubber O-rings (SAKURA SEAL, Tokyo, Japan) or a sham procedure. Subsequently, the mice underwent evaluations for MAC development post-OTAC at intervals of 1, 2, 4, and 8 weeks, alongside assessments of the control group at the outset (Figure 1A). The O-rings had inner diameters of 0.5 and 1.0 mm for primary studies and additional tests, respectively. To compare with the standard transverse aortic constriction (TAC) technique, another set of C57BL/6J male mice weighing 25.0 ± 2.0 g at 9–10 weeks, procured from Oriental Yeast Co., Ltd, Tokyo, Japan, underwent the transverse aortic constriction (TAC) operation[41, 42]. For the generation of SMC lineage tracing mice, a *Myh11-CreER$^{T2}$* mouse and a *ROSA26-EGFP* transgenic mouse were purchased from Jackson Laboratory, and Mhy11-CreER$^{T2}$ mice were crossed with a floxed *ROSA26-EGFP* reporter mouse. *Mhy11-CreER$^{T2}$;ROSA26-EGFP* mice were intraperitoneally injected with 200 mg/10 mL tamoxifen at a dose of 2 mg/10 g body weight for three consecutive days over a week preceding the OTAC procedure. All mice were housed in ventilated cages with unrestricted access to standard rodent diet and purified water and maintained at 22°C with a consistent 12/12 h light/dark cycle.

**Echocardiography**

Transthoracic echocardiography (TTE) was conducted at baseline and weeks 1, 2, 4, and 8 post-OTAC procedure in awake mice using the VEVO 1100 Imaging System (VisualSonics, Toronto, Canada). The procedure has been detailed in a previous study[12].

**Tissue Isolation**

The mice were euthanized via cervical dislocation after sedation with medetomidine, midazolam, and butorphanol at doses of 0.3, 4.0, and 5.0 mg/kg, respectively. Blood was drawn using 23G heparinized needles. To isolate the aorta and heart, the mice were sacrificed by cervical dislocation. The whole heart, followed by the ascending, transverse, and descending aortas, was procured after left ventricular perfusion with phosphate-buffered saline. Next, the surrounding tissues, including adipose tissue, were excised, and the whole heart was harvested to assess cardiac hypertrophy. The aortas were flash-frozen and stored at -80°C or immersed in 10% formalin for subsequent analyses. Finally, the tibial length was measured during autopsy.

**Histological Analysis**

The thoracic aorta was secured and fixed in 10% formalin for at least one day. After embedding in paraffin, the tissues were sectioned at 4 μm. Morphometric evaluations were performed using hematoxylin and eosin-stained aortic sections. Picrosirius red (SR) was used for collagen fiber profiling. Alcian blue was used to identify mucoid extracellular matrix accumulations. Aortic calcification was detected using Von Kossa and Alizarin Red S staining. For the main OTAC experiments, 10 mice per timepoint were analyzed (n = 10), while loose-tie OTAC and standard TAC experiments were conducted 5 mice per group (n = 5). The evaluations included the proximal and distal points of aortic constriction in the OTAC sample and the analogous area in the controls. Images were captured using a fluorescence digital microscope (BZ-X810; Keyence, Osaka, Japan), and aortic wall thickness was analyzed using ImageJ software

(ImageJ 1.53t; National Institution of Health, Montgomery, MD, USA). For quantification of osteochondrogenic cells, H&E sections were evaluated for the presence of chondrocyte-like cells with enlarged, vacuolation and condensed nuclei, consistent with established criteria for hypertrophic chondrocytes[43,44]. The number of osteochondrogenic cells was manually counted per section and normalized to the total number of medial nuclei. Quantification of calcification was performed on von Kossa– and Alizarin Red S–stained sections. The calcified area was measured using ImageJ software and normalized to the total medial area per section.

**Immunohistochemistry and Immunofluorescence**

For immunohistochemical analysis, 4-µm-thick formalin-fixed, paraffin-embedded aortic sections were used. To visualize the localization of alpha-smooth muscle actin (α-SMA), SRY-box transcription factor 9 (SOX9), and RUNX family transcription factor 2 (RUNX2) in tissues, the following procedure was performed. The FFPE aortic sections were deparaffinized using xylene. For antigen retrieval, sections were incubated with 10 mM sodium citrate buffer (pH 6.0, RM102-C, LSI Medience, Tokyo, Japan) for 15 min at 120°C in an autoclave. Endogenous peroxidase blocking was carried out using Dako REAL Peroxidase-Blocking Solution (S202386-2, Dako, Carpinteria, CA, USA) for 10 min at room temperature, and non-specific staining blocking was carried out using

Protein Block Serum-Free (X0909, Dako) for 10 min at room temperature. The slides were then incubated with rabbit anti-α-SMA monoclonal antibody (1:200; #19245, Cell Signaling Technology, Danvers, MA, USA), rabbit anti-SOX9 monoclonal antibody (1:2000; #82630, Cell Signaling Technology), and rabbit anti-RUNX2 monoclonal antibody (1:2000; #12556, Cell Signaling Technology) overnight at 4°C. After washing, the slides were incubated with the appropriate secondary antibodies (EnVision+ System HRP-labelled polymer anti-mouse K4001 or anti-rabbit K4003, Dako) for 30 min at room temperature. The reaction was visualized using 3,3′-diaminobenzidine tetrahydrochloride (DAB) substrate chromogen (K3468, Dako). Finally, the sections were counterstained with hematoxylin (131-09665, FUJIFILM Wako Pure Chemical Corporation, Osaka, Japan) and sealed with a cover glass. The slides were imaged using a fluorescence digital microscope (BZ-X810; Keyence, Osaka, Japan). The proportion of positive staining in aortic tissue was quantified as a percentage using the ImageJ software.

To investigate tissue colocalization, immunofluorescence assays were performed using multiple antibodies. After deparaffinization and antigen retrieval, aortic sections were blocked using Dako REAL Peroxidase-Blocking Solution and Protein Block, Serum-Free, and incubated with the following primary antibody mixtures overnight at 4°C. The primary antibodies used were as follows: mouse anti-α-SMA monoclonal antibody (1:200; ab7817, Abcam Inc., Cambridge, UK), mouse anti-α-SMA monoclonal antibody (1:400;ab270251, Abcam), rabbit anti-SOX9 monoclonal antibody (1:2000; #82630, Cell Signaling Technology), rabbit anti-CD68 monoclonal antibody (1:400; #9778, Cell Signaling Technology), and rabbit anti-Osteopontin/SPP1 (1:400; #88742, Cell Signaling). After washing, the specimens were probed with the following fluorescent secondary antibody mixtures for 60 min at room temperature: Alexa Fluor 488-conjugated goat anti-mouse IgG (1:500; A-11008, Invitrogen, Carlsbad, CA, USA)/cyanine3 goat anti-rabbit IgG [1:500, Invitrogen]) for α-SMA/SOX9. Aortic sections were stained for nuclei with Fluoroshield Mounting Medium using DAPI (ab104139, Abcam) or Hoechst 33258 (H1398, Invitrogen) and observed using an A1 confocal laser microscope (Nikon Co., Tokyo, Japan) or an FSX100 microscope (Olympus, Tokyo, Japan).

**Bulk RNA Sequencing Library Preparation and Sequencing**

Total RNA was extracted from aortic tissues preserved in RNA-later using miRNeasy kits (Cat. No.217084; QIAGEN) and nuclease-free water (Cat. No.9937, Ambion, Austin, TX, USA), according to the manufacturer's protocol (n = 5 per group). RNA quality and quantity were assessed by the A260/280 ratio using a NanoDrop 1000 spectrophotometer (Thermo Fisher Scientific, Wilmington, DE, USA), and the RNA integrity number (RIN) with an Agilent RNA6000 Nano kit (5076-1511, Agilent Technologies, Santa Clara, CA, USA) using an Agilent Bioanalyzer 2100 (G2939A, Agilent Technologies, Santa Barbara, CA, USA). RNA concentration was calculated using a NanoDrop 1000 spectrophotometer. In this study, we set a threshold of RIN > 7.0, indicating excellent RNA quality. Then, poly(A)-selected RNA was used to prepare cDNA libraries with TruSeq Stranded mRNA Library Prep (20020594, Illumina, San Diego, CA, USA) and converted to cDNA. After library construction, single-end sequencing runs (75 cycles) were performed using the NextSeq 500/550 High Output Kit v2.5 (20024906, Illumina) on a NextSeq 500 (Illumina).

**Bulk RNA-seq Data Analyses**

To complement protein-level observations and to address variability in immunostaining, we examined transcript-level dynamics across the OTAC time course using bulk RNA-seq (n = 5 per group). This analysis enabled quantitative assessment of contractile VSMCs markers (e.g., *Acta2*), osteochondrogenic regulators (e.g., *Runx2*), as well as inflammatory and extracellular matrix–related genes, among others. In addition, to compare OTAC aorta with publicly available TAC aorta data, 0-week control and 2-week post-OTAC mice were used. Raw sequencing data were stored in the Sequence Read Archive (NCBI SRA database) under accession number GSE239953. The SRA dataset

(PRJNA659049)[15] was used for comparison with traditional TAC procedures. Raw read quality control and data filtering were performed using fastp (ver. 0.22.0)[45]. Alignments to the reference genome (GRCm39) and quantification of gene expression were conducted using STAR (ver. 2.7.10b)[46] and RSEM (ver. 1.3.3)[47], respectively. Gene expression estimates were imported into the R software (ver. 4.3.2) using the tximport (ver. 1.28.0)[48]. Downstream analysis was conducted using the web-based integrative RNA-seq analysis platform RNAseqChef[49]. Differentially expressed genes (DEGs) between pairwise comparisons were analyzed using the Wald test of DESeq2 with the criteria of fold change (FC) > 2.0 and false discovery rate (FDR) < 0.05. A pathway analysis of significant DEGs was performed using the Gene Ontology (GO) biological process with FDR < 0.05. Gene Set Enrichment Analysis was carried out using the GO biological process on all DEGs.

**scRNA Sequencing Library Preparation and Sequencing**

For the scRNA-seq experiment, we used regions from the ascending aorta to the descending aorta, as this area is expected to represent osteochondrogenic transdifferentiation of vascular smooth muscle cells (VSMCs). To collect time-series data on osteochondrogenic transdifferentiation, 0-week control and 1-, 2-, and 4-week post-OTAC mice were used (Figure 3A). To ensure sufficient cell numbers and biological reproducibility, six mice from each group were pooled. Tissue fixation and dissociation were performed using the Chromium Next GEM Single Cell Fixed RNA Sample Preparation Kit (#1000414,10X Genomics, Pleasanton, CA, USA) according to the manufacturer's protocol. Fixed single cells at 0-week control and 1-, 2-, and 4-week post-OTAC were obtained and subjected to gel bead-in-emulsion (GEMs) construction. Single cells obtained by FACS sorting were subjected to GEM construction to target a single-cell resolution of 10,000 cells. The libraries were constructed using the Chromium Fixed RNA Kit, Mouse Transcriptome (#1000496, 10X Genomics), and sequenced on an Illumina NovaSeq 6000 (Illumina) according to the manufacturer's instructions.

**scRNA-seq Data Analyses**

Raw sequencing data were stored in the NCBI SRA database under accession number GSE268556. Raw FASTQ files from each sample were aligned to the mouse reference genome mm10 using Cell Ranger v7.0.1, and subjected to read filtering, barcode counting, and a unique molecular identifier (UMI) count matrix. The datasets were analyzed using Seurat v5.0.1[50] in R v4.3.2. For quality control, genes expressed in less than 3 cells, cells with > 5% reads mapped to mitochondria, and those with < 200 or > 7,500 UMI counts were removed from downstream analysis. DoubletFinder v2.04[51] was used to remove potential doublets, with the doublet formation rate set to 7.5%

independently for each sample. The doublets were excluded from this study. The SCTransform function with the glmGamPoi[52] method was used for normalization. RunPCA, IntegratedLaysers with Harmony[53], FindNeighbors, FindClusters, and RunUMAP function were also used to reduce dimensionality reduction and remove batch effects. Clustering resolutions ranging from 0.1 to 1.0, with steps of 0.1, were assessed for stability using clustree v0.5.1[54]. Then, the lowest stable resolution of 0.7 was chosen. The cells were then subclustered and annotated based on the manual curation of well-known markers. DEGs were identified using the PrepSCTFindMarkers and FindMarkers functions of Seurat. Biomarkers of different clusters were selected from DEGs with an absolute logFC > 0.25 and FDR < 0.05. To further identify the sub-clusters and annotate them as cell subtypes in each cell type, the previously mentioned steps (normalization, dimensional reduction, and clustering) were repeated for each major cell type. To understand the potential function and mechanism of selected gene lists, such as the marker signature genes of cluster, the enrichment of selected gene list with Gene Ontology (GO) analysis was performed using ClusterProfiler[55] with RNAseqChef[49] with default settings. In addition, pathway activity was assessed at single-cell resolution using AUCell[23] (v1.22.0), which calculates enrichment scores for predefined gene sets in each cell. To systematically infer intercellular communication networks relevant to medial calcification, we applied the LIANA[30] framework (v0.1.14) to our scRNA-seq dataset of the OTAC model. Preprocessed Seurat objects containing human ortholog gene symbols were used as input. Ligand-receptor inference was performed using CellPhoneDB[31] as the primary resource. Clusters annotated as cardiomyocytes were excluded to minimize potential artifacts. For each condition, predicted ligand-receptor pairs were ranked by magnitude mean rank or magnitude rank, depending on availability. To facilitate interpretation, interaction strength was represented as the inverse of that aggregated rank.

**Depletion of Macrophages using Clodronate Liposomes**

Mice were intraperitoneally injected with 625 μg of clodronate liposomes (#16001003, Hygieia Bioscience, Osaka, Japan) in 175 μL of phosphate-buffered saline (PBS) solution to deplete the macrophages (n = 5). For the control group (n = 5), mice were intraperitoneally injected with the same volume of control liposomes (#16003631, Hygieia Bioscience) in 175 μL of PBS solution. Clodronate liposomes or control liposomes were injected one day before the OTAC procedure and every seven days thereafter (Figure 7A). Finally, 28 days after the OTAC procedure, the aortic tissues were harvested for subsequent histological evaluation.

**Neutralization of Ly6g-Positive Cells**

To assess the role of neutrophils in vascular calcification progression in our MAC mouse model, mice were intraperitoneally injected with 352 μg of *InVivo*Plus anti-mouse Ly6G/Ly6C antibodies (#0075, BioXCell, Lebanon, NH, USA) or the *InVivo*Plus rat IgG2b isotype control against keyhole limpet hemocyanin (#BP0090, BioXCell) in 200 μL saline solution (n = 5 in each), as documented in previous studies[56]. The OTAC procedure was initiated two days after the first injection, followed by antibody injections three times per week (Figure 7D). On day 28 post-OTAC, aortic tissues were harvested for subsequent histological evaluation.

**Flow Cytometry**

Peripheral blood was labeled with FITC anti-mouse F4/80 antibody (#123107, clone BM8, BioLegend, San Diego, CA, USA) and PE anti-mouse CD11c antibody (#117307, N418, BioLegend) for assessment of macrophages or Alexa Fluor® 488 anti-mouse/human CD11b antibody (#101219, M1/70, BioLegend) and APC anti-mouse Ly6G antibody (#127613, 1A8, BioLegend) for assessment of neutrophils. Peripheral blood samples with the antibodies were lysed using OptiLyse C Lysing Solution (A11895; BD Biosciences, San Jose, CA, USA). After lysis, the samples were rinsed with a solution containing 10% fetal bovine serum to remove the residual lysis solution and cellular debris. Analyses were performed using a FACSCalibur flow cytometer (BD Biosciences).

**Statistical Analyses**

Data are presented as the mean ± standard error of the mean or median and interquartile range. Statistical analyses were performed using GraphPad Prism ver. 10.0.1 (GraphPad Software, San Diego, CA, USA), deploying either an unpaired t-test or one-way analysis of variance with Tukey's post hoc test. Statistical significance was set at $P < 0.05$.

**Code Availability**

The source code used to reproduce our analysis can be accessed upon reasonable request from the corresponding authors.

### Supplemental Figures

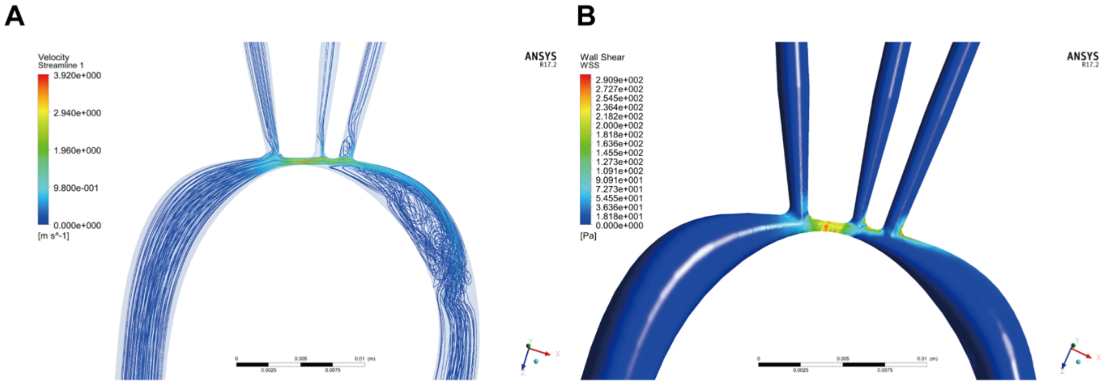


**Supplemental Figure S1. Simulation of blood flow velocity and wall shear stress in OTAC mouse aorta using ANSYS fluent**

**A**: Streamlines simulated for the aorta in O-ring-induced transverse aortic constriction (OTAC) mice. Increased blood flow velocity at the stenosis site was observed. **B**: Wall shear stress (WSS) simulated in the aorta of OTAC mice. A WSS peak was observed at the stenotic site.

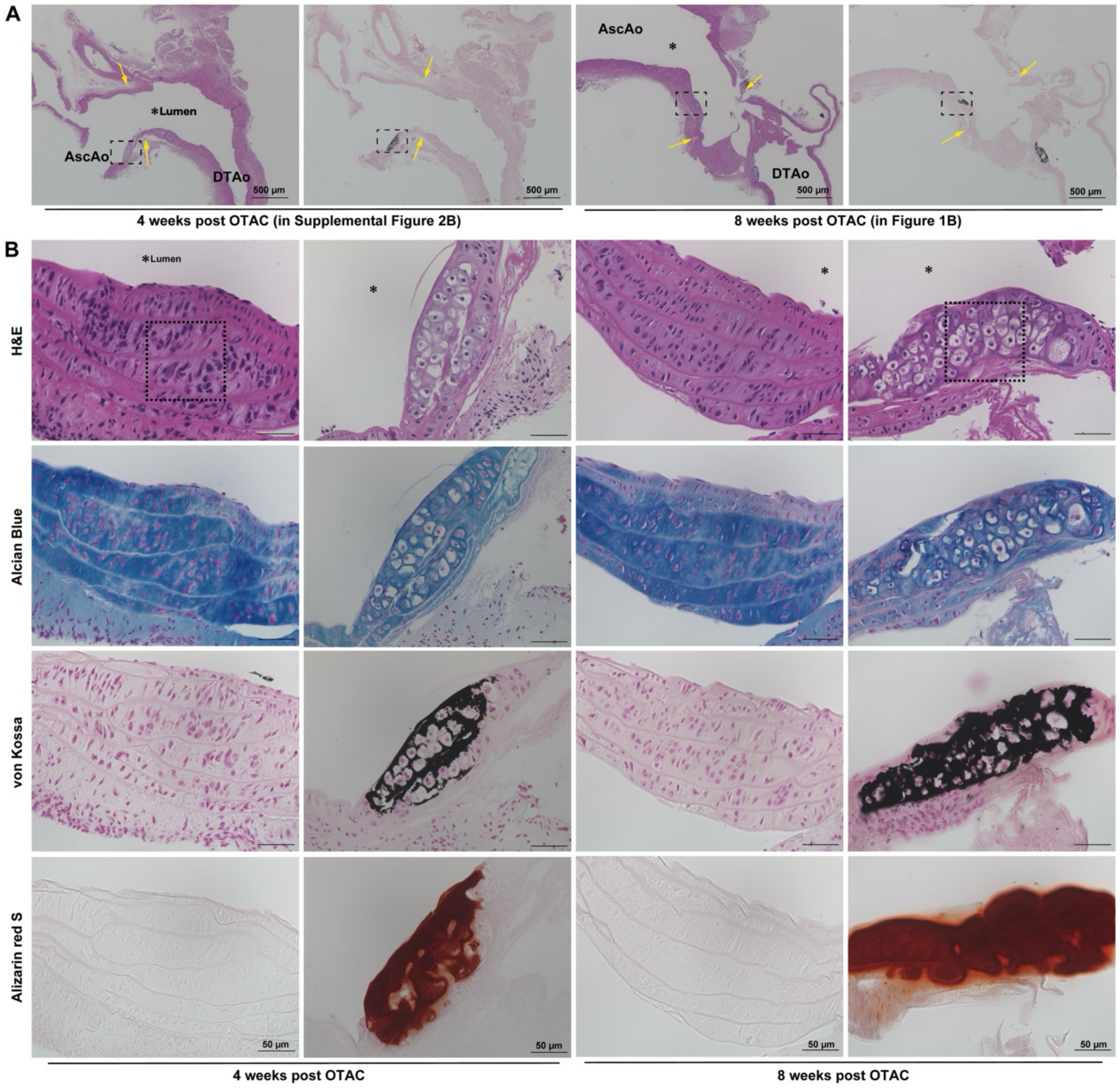


**Supplemental Figure S2. Representative histological images from multiple animals at 4- and 8-week post-OTAC**

**A.** Longitudinal sections of the aorta at 4 and 8 weeks post-OTAC (left, H&E; right, von Kossa), shown at ×40 magnification. The 4-week panel represents the same specimen as shown in Supplemental Figure 2B (second column from the left). The 8-week panel corresponds to the lower-magnification image presented in main Figure 1B. Yellow arrows indicate the OTAC-induced stenosis region. **B.** Higher-magnification transverse sections (×400) from 4- and 8-week post-OTAC specimens stained with H&E, Alcian blue, von Kossa, and Alizarin Red S.

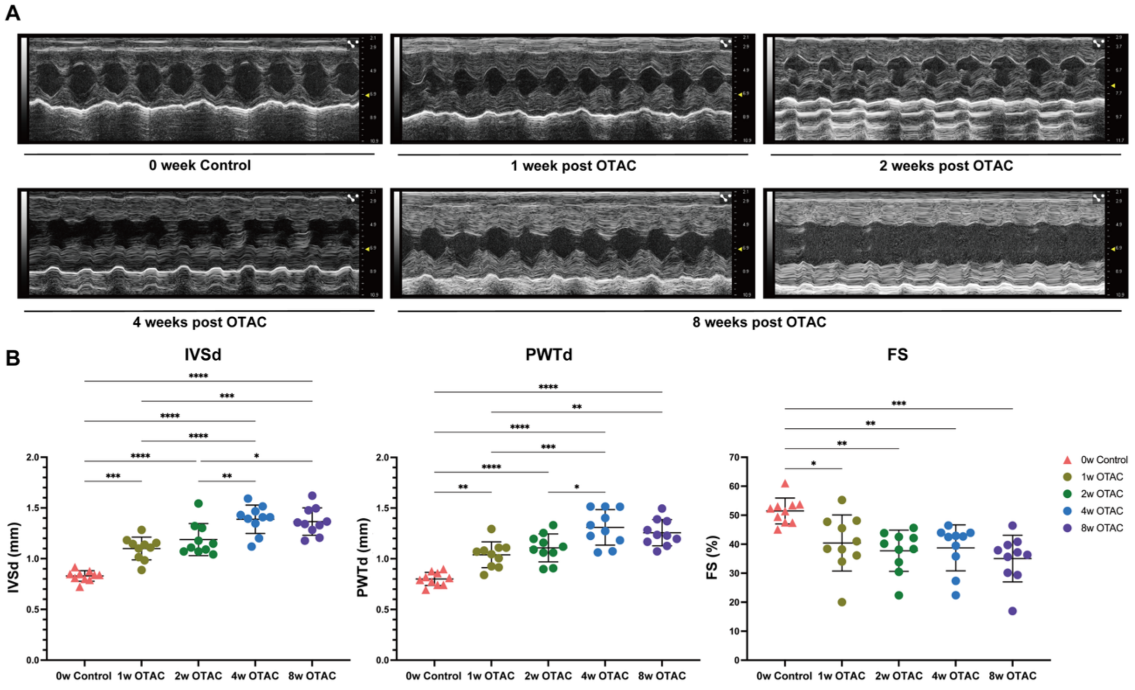


**Supplemental Figure S3. Echocardiographic analyses before and after OTAC and Sham.**

**A.** Representative echocardiographic M-mode of mice 1-, 2-, 4-, and 8-week post-OTAC procedure and 0-week control mice. The x-axis represents the time, and the y-axis represents the distance (in mm) from the transducer. **B.** Echocardiographic measurements of interventricular septum in diastole: IVSd, left ventricular posterior wall thickness in diastole: PWTd, and LV fractional shortening: FS. Data are presented as mean ± standard error of the mean. Group comparisons were performed using one-way analysis of variance with Tukey's post-hoc test. Significance levels are denoted as *P < 0.05; **P < 0.01; ***P < 0.001; ****P < 0.0001.

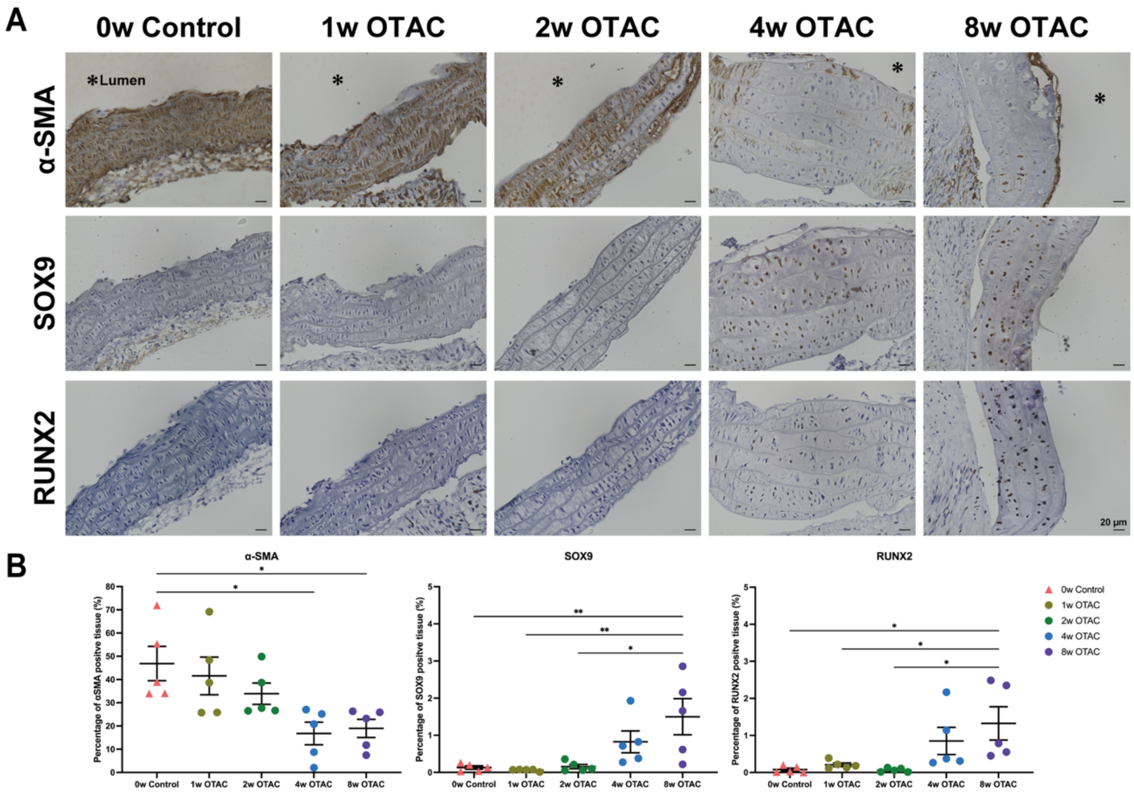


**Supplemental Figure S4. Transition of vascular smooth muscle cells from α-SMA-positive to RUNX2- and SOX9-positive cells.**

**A.** Immunohistochemical analysis of α-SMA, SOX9, and RUNX2 in the aortas of mice at 1-, 2-, 4-, and 8-week post-O-ring-induced transverse aortic constriction (OTAC) compared to controls. Scale bar, 20 µm. **B**. Quantitative analysis of the tissue areas positive for α-SMA, SOX9, and RUNX2 relative to the total tissue area. Data are presented as mean ± standard error of the mean. Group comparisons were conducted using one-way analysis of variance, followed by Tukey's post-hoc test. $^*P < 0.05$; $^{**}P < 0.01$.

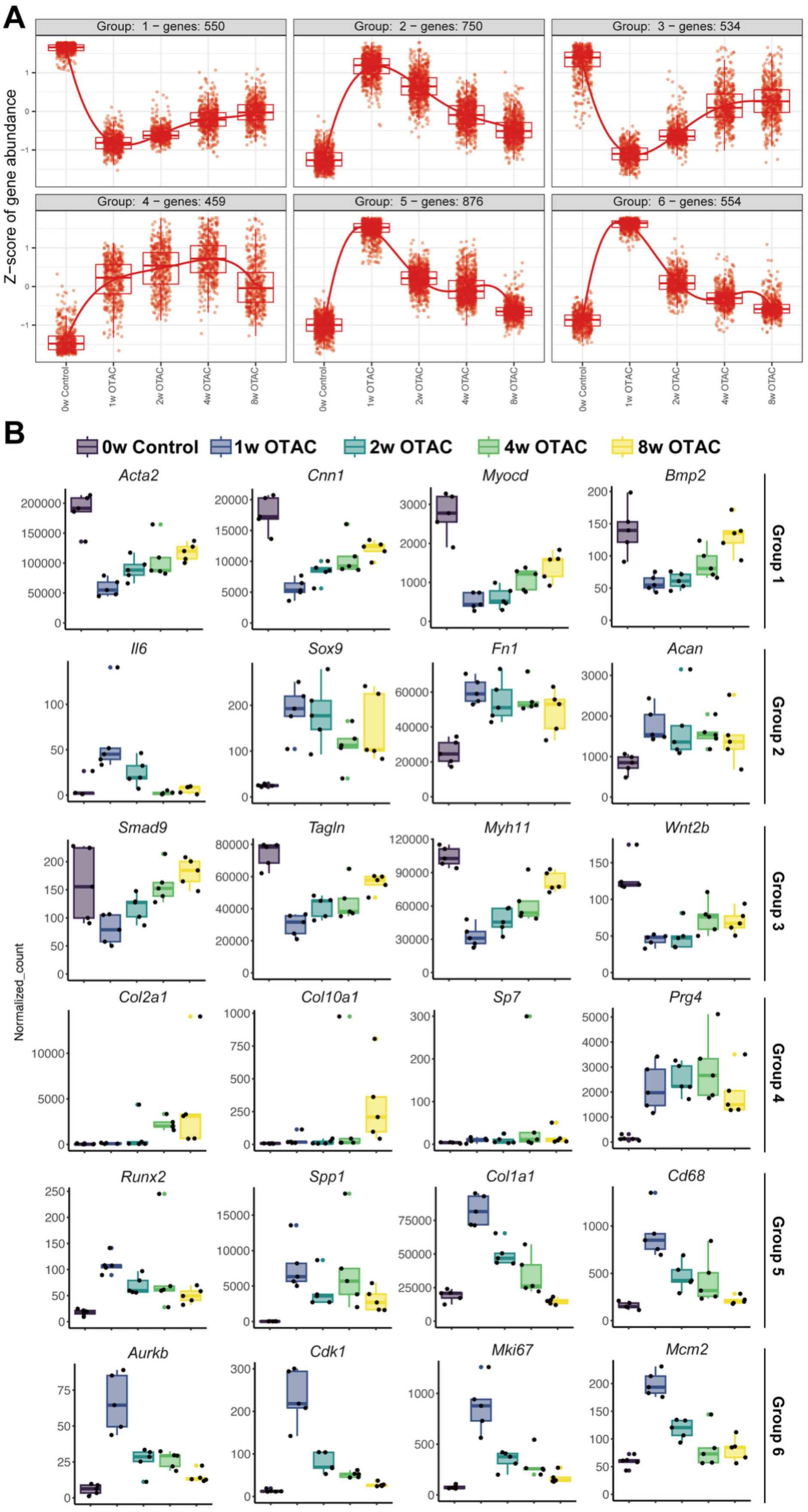
A
Group: 1 – genes: 550
Group: 2 – genes: 750
Group: 3 – genes: 534
Group: 4 – genes: 459
Group: 5 – genes: 876
Group: 6 – genes: 554
Z−score of gene abundance
0w Control
1w OTAC
2w OTAC
4w OTAC
8w OTAC
B
0w Control
1w OTAC
2w OTAC
4w OTAC
8w OTAC
Acta2
Cnn1
Myocd
Bmp2
Group 1
Il6
Sox9
Fn1
Acan
Group 2
Smad9
Tagln
Myh11
Wnt2b
Group 3
Normalized_count
Col2a1
Col10a1
Sp7
Prg4
Group 4
Runx2
Spp1
Col1a1
Cd68
Group 5
Aurkb
Cdk1
Mki67
Mcm2
Group 6

**Supplemental Figure S5. Transcript-level dynamics during OTAC-induced calcification.**

Bulk RNA-seq analysis (n = 5 per group) was performed across the OTAC time course. **A.** K-means clustering stratified transcripts into six groups. **B.** Representative temporal gene expression changes in each group. OTAC, O-ring-induced transverse aortic constriction; w, week.

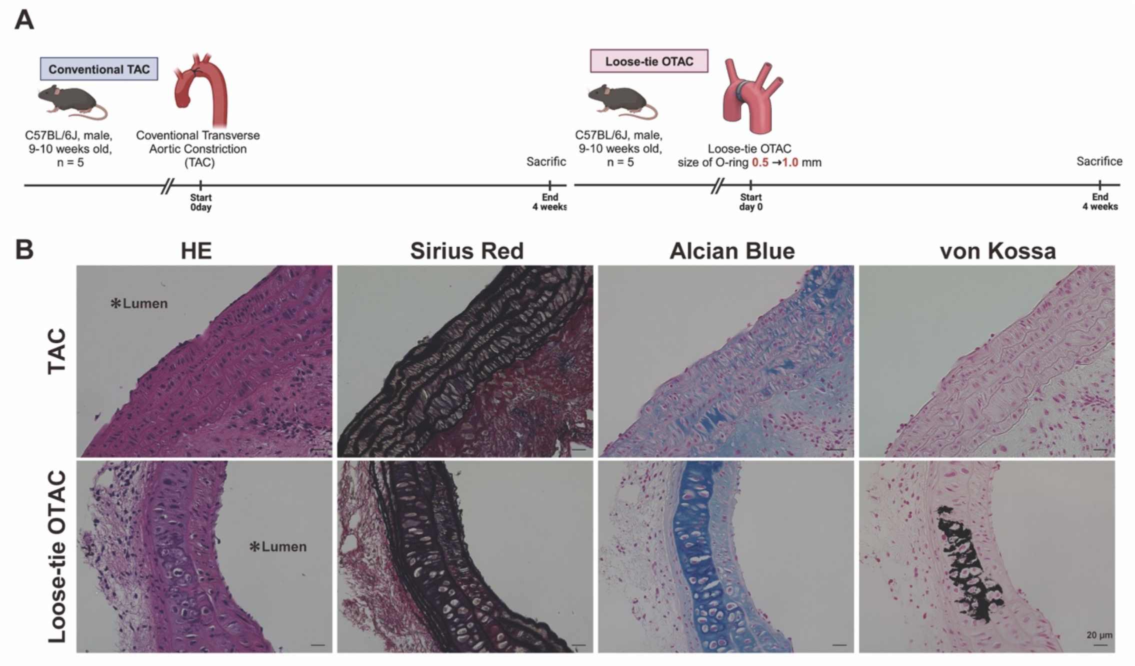


**Supplemental Figure S6. Aortic histological analysis in the TAC and loose-tie OTAC mice.**

**A.** Conventional transverse aortic constriction (TAC) and loose-tie O-ring-induced aortic constriction (OTAC) examination. **B.** Distinctive aortic tissue histological analyses were evaluated with hematoxylin and eosin (HE), Sirius Red, Alcian Blue, and von Kossa for the TAC and loose-tie OTAC groups. A scale bar is indicated in each image and corresponds to 20 μm. For histological evaluation, five mice were considered per group.

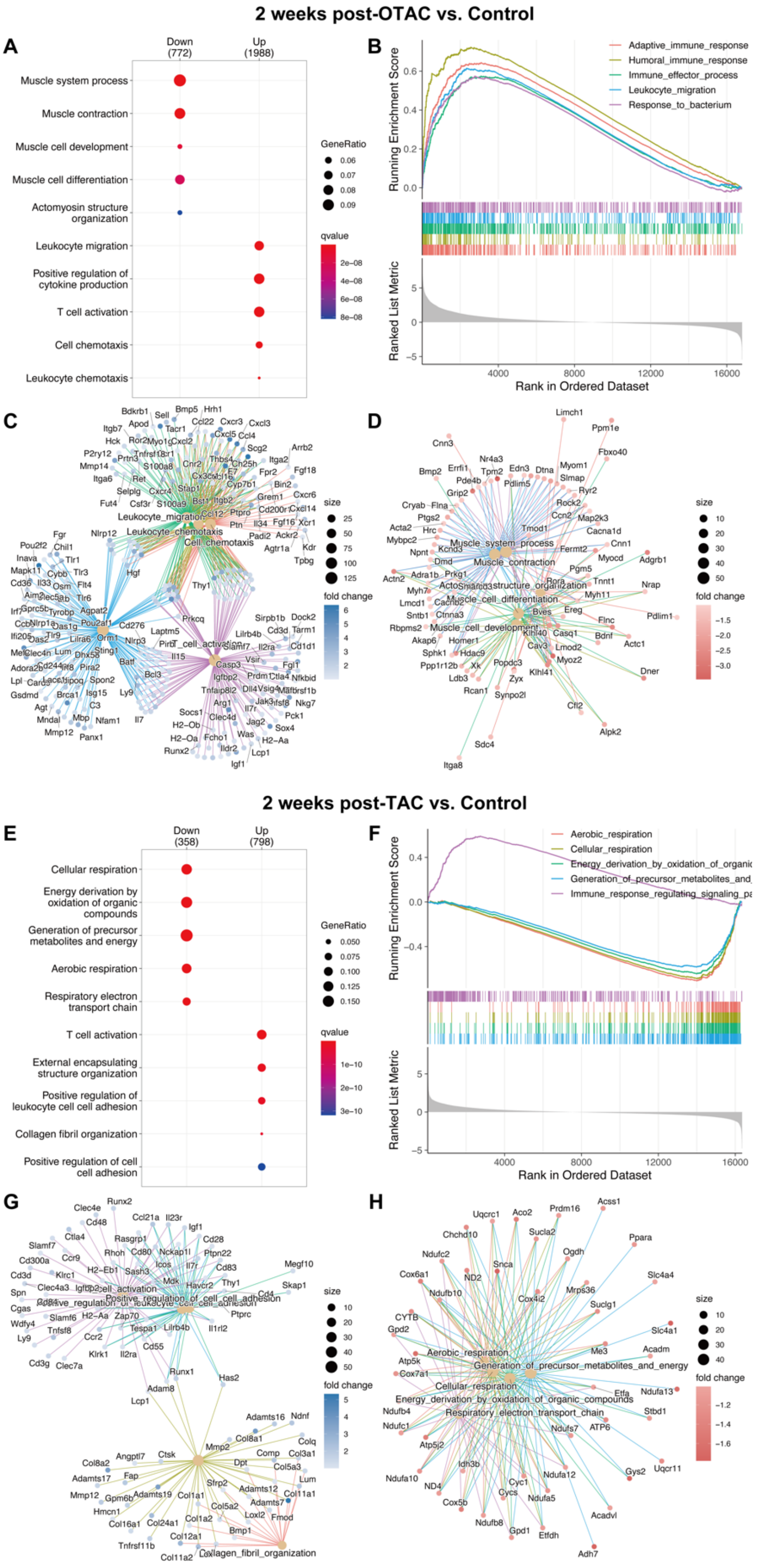
2 weeks post-OTAC vs. Control
A
Down (772)
Up (1988)
Muscle system process
Muscle contraction
Muscle cell development
Muscle cell differentiation
Actomyosin structure organization
Leukocyte migration
Positive regulation of cytokine production
T cell activation
Cell chemotaxis
Leukocyte chemotaxis
GeneRatio
qvalue
B
Running Enrichment Score
Ranked List Metric
Rank in Ordered Dataset
Adaptive_immune_response
Humoral_immune_response
Immune_effector_process
Leukocyte_migration
Response_to_bacterium
C
size
fold change
D
2 weeks post-TAC vs. Control
E
Down (358)
Up (798)
Cellular respiration
Energy derivation by oxidation of organic compounds
Generation of precursor metabolites and energy
Aerobic respiration
Respiratory electron transport chain
T cell activation
External encapsulating structure organization
Positive regulation of leukocyte cell cell adhesion
Collagen fibril organization
Positive regulation of cell cell adhesion
F
Aerobic_respiration
Cellular_respiration
Energy_derivation_by_oxidation_of_organic
Generation_of_precursor_metabolites_and
Immune_response_regulating_signaling_pa
G
H

**Supplemental Figure S7. Gene ontology biological processes and gene set enrichment analysis of aortic tissue in O-ring-induced transverse aortic constriction and transverse aortic constriction mice.**

Analysis of the up- and downregulated differentially expressed genes (DEGs) derived from two datasets: **A-D.** Comparison between 2-week post-OTAC and 0-week control mice. **E-H.** Comparison between 2-week post-conventional TAC mice from the PRJNA65904934 dataset and their corresponding control mice. **A and E.** The top five signaling pathways based on the Gene Ontology (GO) biological processes were enriched in gene sets, including 1,988 upregulated and 772 downregulated genes between the OTAC and control mice (**A**) and 798 upregulated and 358 downregulated genes between the TAC and control mice (**E**) with a threshold of fold change (FC) > 2.0, and false discovery rate (FDR) < 0.05. **B and F.** Gene set enrichment analysis (GSEA): Top five gene set enrichment score plots for the indicated gene sets comparing the OTAC and control mice (**B**) and the TAC and control mice (**F**). **C-D and G-H.** Gene-concept networks: The top five upregulated GO terms in OTAC (**C)** and TAC mice (**G**) and downregulated GO terms in OTAC (**D**) and TAC mice (**H**). OTAC, O-ring-induced transverse aortic constriction; TAC, transverse aortic constriction.

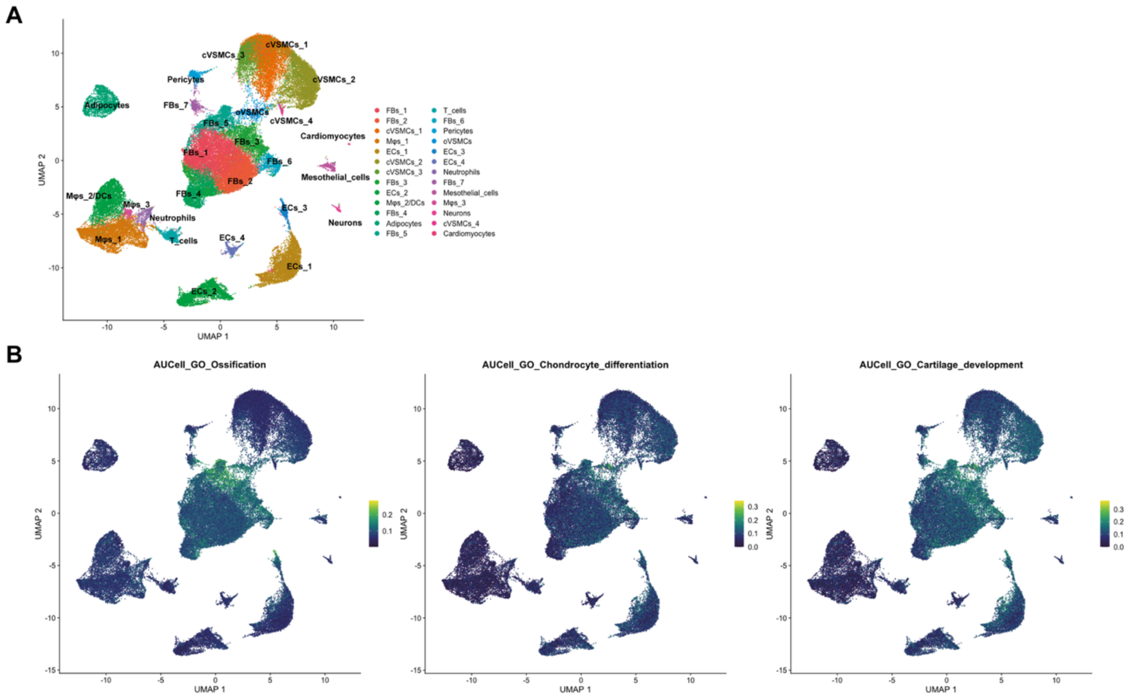


**Supplemental Figure S8: AUCell analysis of pathway activity at single-cell resolution.**

**A.** Uniform Manifold Approximation and Projection (UMAP) plots of all cells, color-coded by cell type. **B.** UMAP feature plots show per-cell enrichment scores calculated by AUCell for representative Gene Ontology biological process terms, including ossification, chondrocyte differentiation, and cartilage development.

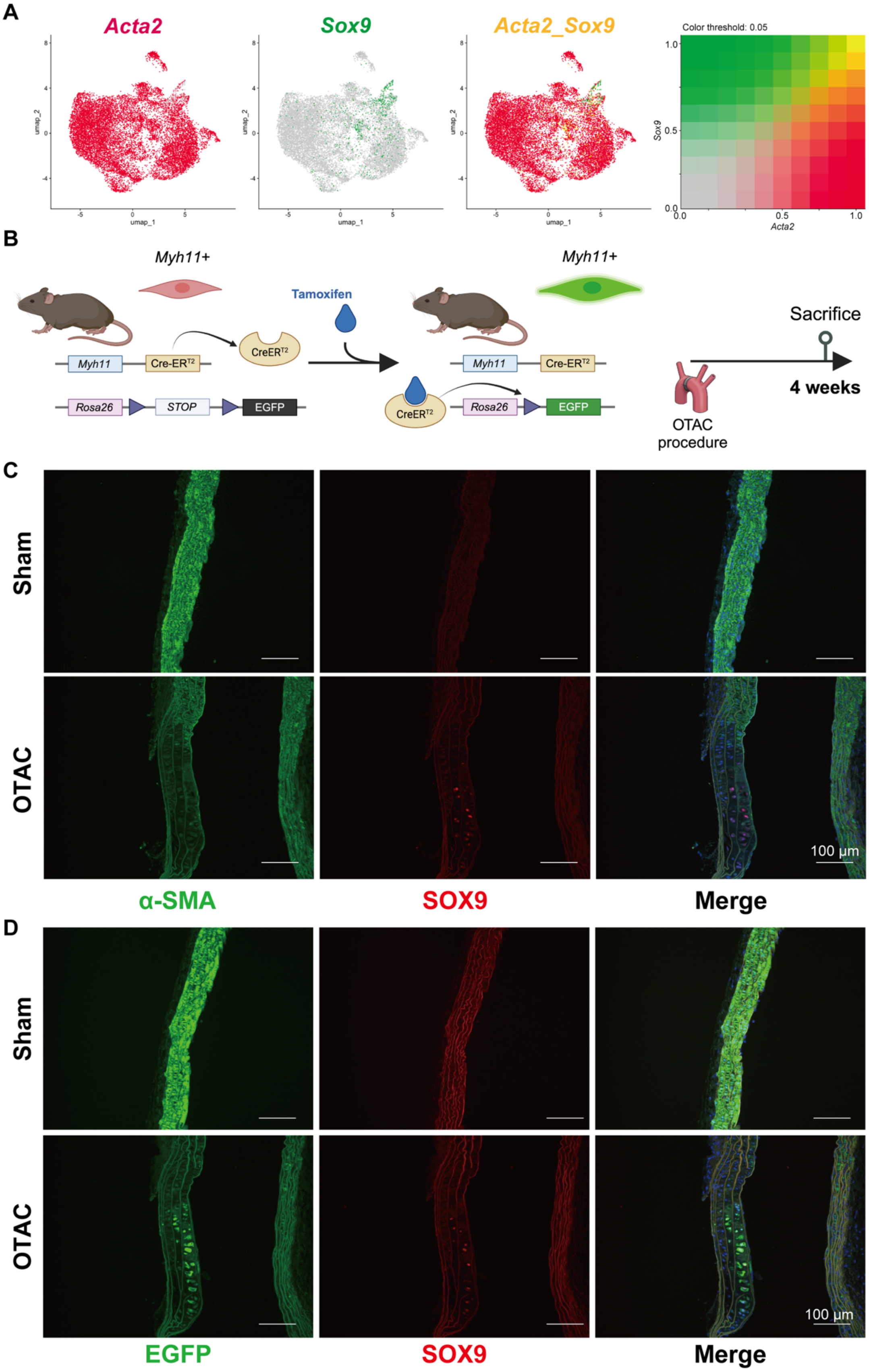
A
Acta2
Sox9
Acta2_Sox9
Color threshold: 0.05
umap_1
umap_2
Sox9
Acta2
B
Myh11+
Tamoxifen
Myh11
Cre-ERT2
CreERT2
Rosa26
STOP
EGFP
Sacrifice
4 weeks
OTAC procedure
C
Sham
OTAC
100 μm
α-SMA
SOX9
Merge
D
Sham
OTAC
100 μm
EGFP
SOX9
Merge

**Supplemental Figure S9**. **Lineage tracing analysis revealed α-SMA and SOX9 double-positive cells derived from MYH11-positive cells.**

**A.** UMAP plots shows the relative expression of *Acta2* (**left**), *Sox9* (**middle**), and both of them (**right**) in the VSMCs sub-cluster. **B.** Study overview. *Myh11Cre-ER*$^{T2}$;*ROSA26-EGFP* mice were used for the lineage tracing. **C-D.** Immunohistochemical analysis of α-SMA and SOX9 (**C**) and EGFP and SOX9 (**D**) in *Myh11-CreER*$^{T2}$;*ROSA26-EGFP* mice. For both panels, upper rows show sham-operated controls and lower rows show OTAC-operated aortas. Scale bar: 100 μm. OTAC, O-ring-induced transverse aortic constriction.

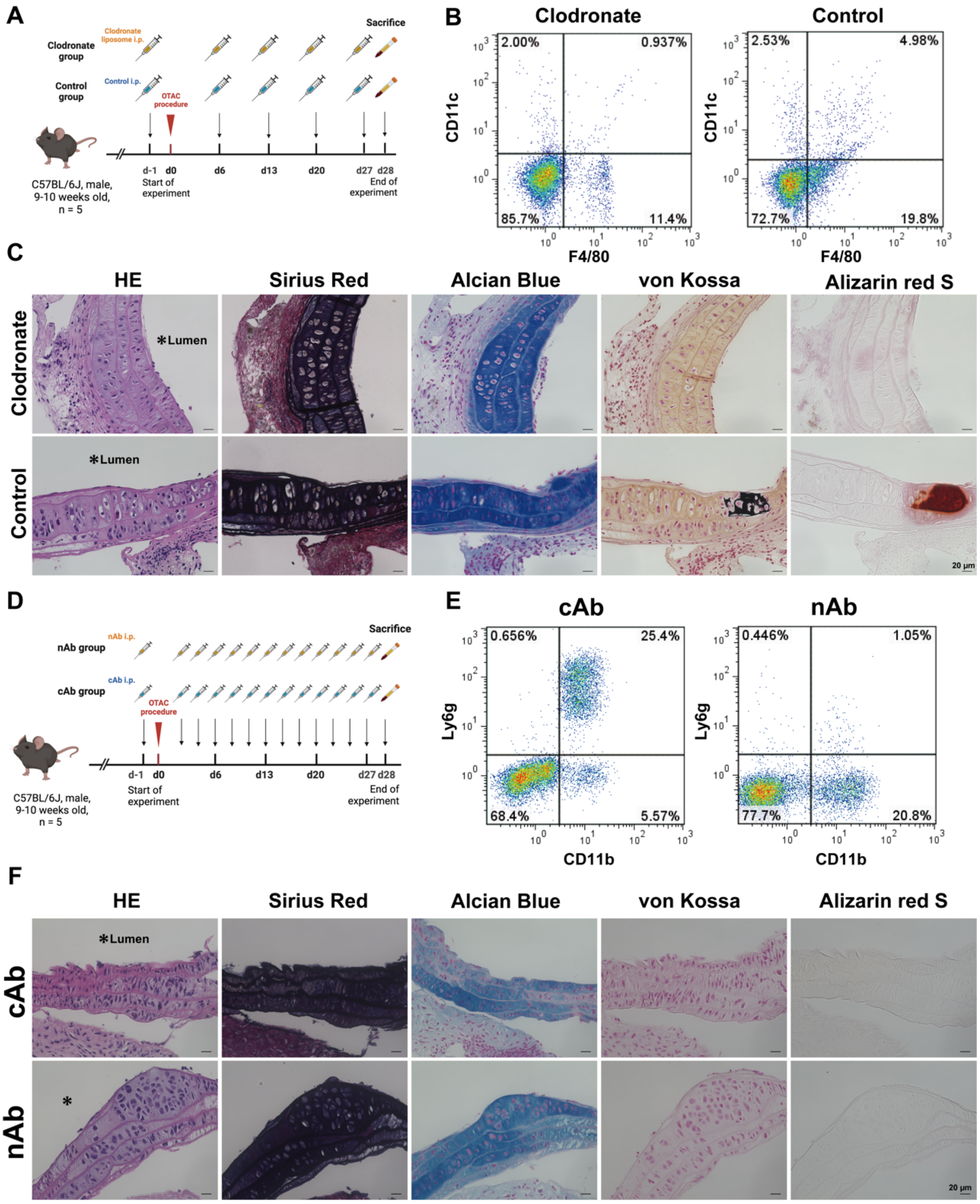


**Supplemental Figure S10: Injection of Clodronate Liposome and Neutralization of Neutrophils for O-ring-induced transverse aortic constriction mice.**

**A.** Administration protocol for clodronate liposomes and control liposomes. **B.**

Representative flow cytometry analysis of F4/80-positive and CD11c-positive cells in peripheral blood from O-ring-induced transverse aortic constriction (OTAC) mice treated with clodronate or control liposomes (day 28). **C.** Representative aortic tissue images of hematoxylin and eosin (HE), Sirius Red, Alcian Blue, and von Kossa staining in each group. **D.** Protocol for Administration of anti-Ly6g neutralizing antibodies (nAb) or isotype control antibodies (cAb). **E.** Representative flow cytometry analysis of Ly6g-positive and CD11b-positive cells in peripheral blood from O-ring-induced transverse aortic constriction (OTAC) mice treated with nAb or cAb (day 28). **F.** Representative aortic tissue images of hematoxylin and eosin (HE), Sirius red, Alcian blue, and von Kossa staining in each group. Scale bar: 20 μm. HE, hematoxylin and eosin; OTAC, O-ring-induced transverse aortic constriction.

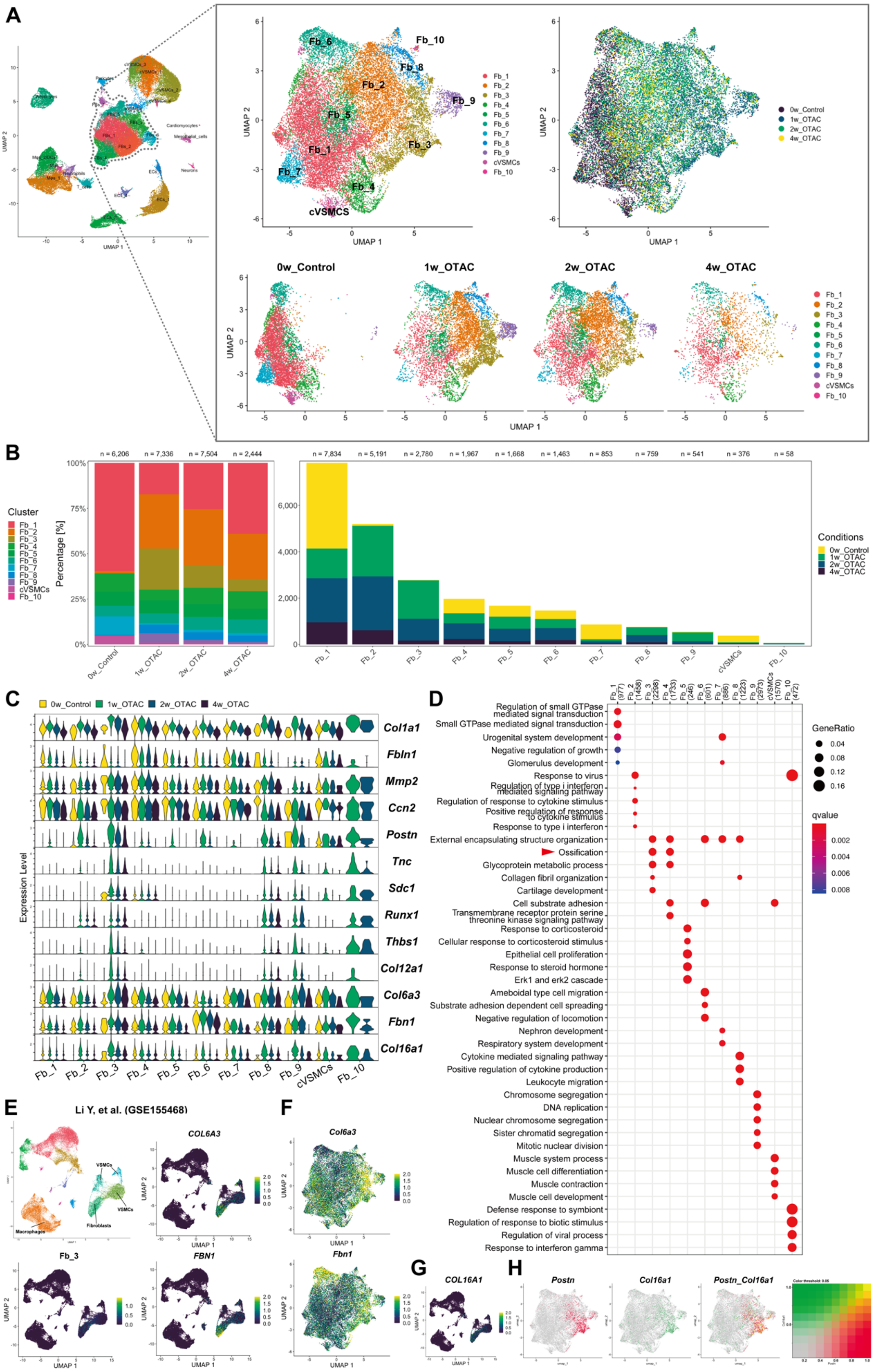
A
Fb_1
Fb_2
Fb_3
Fb_4
Fb_5
Fb_6
Fb_7
Fb_8
Fb_9
Fb_10
cVSMCS
cVSMCs
0w_Control
1w_OTAC
2w_OTAC
4w_OTAC
UMAP 1
UMAP 2
B
Cluster
Percentage [%]
Conditions
n = 6,206
n = 7,336
n = 7,504
n = 2,444
n = 7,834
n = 5,191
n = 2,780
n = 1,967
n = 1,668
n = 1,463
n = 853
n = 759
n = 541
n = 376
n = 58
C
Expression Level
Col1a1
Fbln1
Mmp2
Ccn2
Postn
Tnc
Sdc1
Runx1
Thbs1
Col12a1
Col6a3
Fbn1
Col16a1
D
GeneRatio
qvalue
Regulation of small GTPase mediated signal transduction
Small GTPase mediated signal transduction
Urogenital system development
Negative regulation of growth
Glomerulus development
Response to virus
Regulation of type i interferon mediated signaling pathway
Regulation of response to cytokine stimulus
Positive regulation of response to cytokine stimulus
Response to type i interferon
External encapsulating structure organization
Ossification
Glycoprotein metabolic process
Collagen fibril organization
Cartilage development
Cell substrate adhesion
Transmembrane receptor protein serine threonine kinase signaling pathway
Response to corticosteroid
Cellular response to corticosteroid stimulus
Epithelial cell proliferation
Response to steroid hormone
Erk1 and erk2 cascade
Ameboidal type cell migration
Substrate adhesion dependent cell spreading
Negative regulation of locomotion
Nephron development
Respiratory system development
Cytokine mediated signaling pathway
Positive regulation of cytokine production
Leukocyte migration
Chromosome segregation
DNA replication
Nuclear chromosome segregation
Sister chromatid segregation
Mitotic nuclear division
Muscle system process
Muscle cell differentiation
Muscle contraction
Muscle cell development
Defense response to symbiont
Regulation of response to biotic stimulus
Regulation of viral process
Response to interferon gamma
E
Li Y, et al. (GSE155468)
COL6A3
FBN1
Fb_3
VSMCs
Macrophages
Fibroblasts
F
Col6a3
Fbn1
G
COL16A1
H
Postn
Col16a1
Postn_Col16a1

**Supplemental Figure 11. Characterization of fibroblast subpopulations in OTAC and control mice.**

**A.** Uniform Manifold Approximation and Projection (UMAP) plot of all fibroblasts, colored according to fibroblasts sub-clusters (**upper left**), the originating group (**upper right**), and divided by the originating group (**lower**) from 0-week control (0w_Control), 1-week post-O-ring-induced transverse aortic constriction (1w_OTAC), 2-week post-OTAC (2w_OTAC), and 4-week post-OTAC (4w_OTAC). Cell numbers for each group are as follows: 0w_Control (n = 6,206 cells from six mice), 1w_OTAC (n = 7,336 cells from six mice), 2w_OTAC (n = 7,504 cells from six mice), and 4w_OTAC (n = 2,444 cells from six mice). **B.** Proportion of cell types in each group (**left**). Number of cells in each sub-cluster, colored by group (**right**). **C.** Violin plots showing fibroblast-specific markers, colored for each group. **D.** Top five enriched Gene Ontology (GO) biological process terms of macrophage sub-clusters. Red and blue arrowheads indicate GO terms directly associated with vascular calcification. **E.** UMAP visualization from Li et al. single-cell RNA-sequencing (scRNA-seq) data (n = 45,754 cells from eight patients with ascending thoracic aortic aneurysm and three controls) showing *COL6A3, FBN1,* and Fb_3 modulation score. **F.** UMAP plot colored by the relative expression of *Col6a3* and *Fbn1*. **G.** UMAP visualization from Li et al. scRNA-seq data showing *COL16A1* expression. **H.** UMAP shows the relative expression of *Postn* (**left**), *Col16a1* (**middle**), and both (**right**) in the fibroblast sub-cluster. VSMC, vascular smooth muscle cell; cVSMC, contractile VSMC; Fb, fibroblast

**Supplemental Table**

**Supplemental Table S1. Summary of cell numbers before and after quality control for scRNA-seq in this study**

| **Sample** | **Cellranger output** | **After quality control** | **Removing doublets** | **Remaining singlets** |
|---|---|---|---|---|
| 0w_Control | 18,038 | 18,015 | 1,014 | 17,001 |
| 1w_OTAC | 22,831 | 22,662 | 1,309 | 21,353 |
| 2w_OTAC | 19,895 | 19,821 | 1,140 | 18,681 |
| 4w_OTAC | 6,062 | 6,048 | 340 | 5,708 |
| **Total cell numbers** | 66,826 | 66,546 | 3,803 | 62,743 |

Note: Cells expressing > 200 and < 7500 genes, as well as genes expressed in more than three cells were selected. Cells with ≥ 5% mitochondria-derived genes were screened. After removing doublets from each sample, 62,743 cells were used for subsequent analyses.
